# Aberrant immune regulation and enrichment of stem-like CD8^+^ T cells in the pancreatic lymph node during type 1 diabetes development

**DOI:** 10.1101/2025.05.23.655848

**Authors:** Leeana D. Peters, Howard R. Seay, Justin Smith, Amanda L. Posgai, Reed Berkowitz, Clive H. Wasserfall, Mark A. Atkinson, Rhonda Bacher, Maigan A. Brusko, Todd M. Brusko

**Author notes:** Correspondence to: Todd M. Brusko, Ph.D., Department of, College of Medicine, University of Florida, 1275 Center Drive, Biomedical Sciences Building J589, Box 100275, Gainesville, FL 32610.

## Abstract

Effector CD8^+^ T cells are key cellular drivers of type 1 diabetes (T1D) pathogenesis, yet questions remain regarding the molecular defects leading to altered cytotoxicity, their signature in peripheral tissues, and their receptor specificity. Thus, we analyzed human pancreatic lymph nodes (pLN) using mass cytometry and single cell RNA sequencing (scRNAseq) with combined proteomic and T cell receptor (TCR) profiling. Cytometric analysis revealed an enriched population of T stem-cell memory (TSCM)-like cells (CD8^+^CD45RA^+^CD27^+^CD28^+^CCR7^+^CXCR3^+^ T cells) in T1D pLNs. scRNAseq profiling indicated an elevated inflammatory cytokine gene signature (*IFITM3*, *LTB*) along with regulators of terminal differentiation (*BCL6*, *BCL3*), coupled with reduced expression of exhaustion-associated genes (*DUSP2*, *NR4A2*, *TSC22D3*) in CD8^+^ T cells in T1D pLN. Additionally, effector CD8^+^ T cells expressed features of progenitor exhausted cells (*BCL2*) in T1D pLN. Immune Response Enrichment Analysis (IREA) indicated IL-15 signaling as a significant driver of these phenotypes. Integrated TCR and transcriptomic analysis revealed a cluster of diverse naïve-like CD8^+^ T cell clones in T1D pLN. When comparing pLN and pancreatic slice cellular isolates, we observed sharing of effector CD8^+^ T cells, with upregulation of terminal effector signatures detected within the pancreas relative to paired pLN samples. Multiplex imaging revealed differential localization of TCF1 and TOX expressing T cells in the pancreas, with TCF1^+^TOX^+^ cells located in closer proximity to the islets and displaying a mixture of activation and exhaustion-associated phenotypes. Thus, we provide multimodal cellular profiles enriched in T1D tissues for consideration in therapeutic targeting.

## Introduction

Genetic fine mapping studies have identified that type 1 diabetes (T1D)-associated risk variants preferentially localize to enhancer regions within immune cells, namely T and B cells^1^. However, inaccessibility of the pancreas from living individuals mandates that studies examining immune cell function in T1D largely be derived from peripheral blood, which may not accurately reflect phenotypes within the target organ during disease pathogenesis. Identifying the molecular basis for T1D-associated immune dysregulation within the pancreas and pancreatic draining lymph nodes (pLN) is highly valuable in the process of developing targeted therapies.

The role of the pLN in promoting T1D progression is incompletely characterized, but murine and human studies have implicated the pLN as an important priming site for immune activation^2–4^. In the non-obese diabetic (NOD) mouse model of T1D, pLN self-renewing autoreactive T cells were found to be an important source of islet antigen-specific cytotoxic lymphocytes^5^. Human studies indicate that patients with T1D possess skewed T cell subset distributions within the pLN, with reduced T follicular regulatory (TFR) cell frequency reported in one study^6^ and increased Th17 cells noted in another^7^. Although there are conflicting reports as to whether deficits in circulating regulatory T cell (Treg) number and function are present in T1D patients^8, 9^, multiple studies have demonstrated deficits in suppressive capacity^7^ and migration toward the pancreas^10^ in Tregs derived from T1D pLN.

Though peripheral blood signatures do not always correlate with those from tissues^11^, there have been many important knowledge gains linking peripheral immunological signatures to clinical outcomes. Interestingly, many of the signatures associated with immunotherapeutic response have converged around CD8^+^ T cell subsets. For example, an exhausted-like CD8^+^ population expressing negative regulatory molecules, including TIGIT and PD-1, together with exhaustion markers KLRG1 and CD57, was associated with C-peptide preservation post-treatment with alefacept (LFA3-Ig)^12^. Similarly, expansion of TIGIT^+^KLRG1^+^CD8^+^ T cells was found preferentially in those who responded to teplizumab (α-CD3)^13^.

Moreover, studies in mice^5^ and humans^14, 15^ have implicated CD4^+^ and CD8^+^ stem cell memory T cells (TSCM) as a potentially important reservoir for promoting pathogenic antigen-specific responses in T1D. Although clinical studies have shown that exhausted-like populations are clonally related to a different CD8^+^ memory population expressing more activation markers relative to exhaustion markers^12^, there is a paucity of information in T1D regarding the distribution of these subsets in secondary lymphatic organs and pancreas, their receptor profiles, or the pathways driving these phenotypes.

Our lab has previously generated sorted T cell receptor (TCR) profiles from donor-matched pLN, spleen, and peripheral blood samples, revealing cell type-dependent repertoire sharing across tissues, with CD8^+^ T cells demonstrating the greatest degree of sharing^16^. The advent of single cell profiling technologies enables the examination of sharing between lymph node and pancreas T cell populations with greater resolution. To understand the phenotypic and transcriptomic profile of immune cell populations in disease-relevant tissues, we performed single cell multi-omic analysis of the pLN as well as pancreatic T cells from donors with and without T1D. In this study, we show that circulating phenotypic signatures previously identified to correlate with T1D status or immunotherapeutic response are shared across tissue immune phenotypes. Moreover, we offer evidence that human CD8^+^ T cells possess fewer differentiated phenotypes in the pLN in T1D and that effector populations differentiate further upon trafficking to the pancreas.

## Methods

### Sex as a biological variable

Male and female organ donors were accepted in this study.

### Human Organ Donors

Tissues were recovered from T1D donors, donors possessing islet autoantibody positive (AAB+), as well as donors without diabetes and negative for islet autoantibodies (ND) according to the Network for Pancreatic Organ donors with Diabetes (nPOD) inclusion criteria^17^. Demographic information used to stratify subjects for analyses was obtained from the nPOD datashare (https://npoddatashare.coh.org/) and nPOD data portal (https://portal.jdrfnpod.org) websites. See **Table S1** for a description of cases and corresponding experiment information.

### Organ Processing

Tissues were processed as previously described^16^. Briefly, pLN were dissociated in HBSS + 10% human (male, type AB) serum (Atlanta Biologicals) using a gentleMACS tissue processor (Miltenyi), followed by subsequent homogenization and filtering steps using a tissue douncer (40 mL; loose pestle; Wheaton) and 100-μm, 70-μm, and 30-μm filters (Miltenyi) to obtain a single cell suspension. Cells were freshly analyzed by flow cytometry as described below, and remaining cells were cryopreserved using CryoStor CS10 (Stemcell). Live pancreatic slices were prepared from nPOD organ donors as previously described^18^ and freshly digested with 1X collagenase IV (Stemcell) diluted in DMEM for 30 minutes at 37°C and 5% CO_2_. Remaining tissue was gently mechanically dissociated with frosted slides and filtered using 70-μm filters (Miltenyi) to obtain a single cell suspension. Cells were enriched for CD45^+^ cells using CD45 Microbeads (Milltenyi) and processed for scRNAseq (10X Genomics) as below.

### Flow Cytometry

Fresh tissues were dissociated as above and stained with Live/Dead yellow viability dye (Invitrogen) and incubated with TruStain FcX (Biolegend) according to the manufacturer’s instructions prior to staining with a panel of antibodies broadly examining memory, effector, and naïve immune cell phenotypes (**Table S2**) consistent with previously published methods^19^. Data were acquired on an LSRFortessa (BD). To assess expression of transcription factors TCF1 and TOX in CD8^+^ T cell populations, cryopreserved pLN cell suspensions were thawed, stained with Zombie Aqua viability dye (Invitrogen), incubated with TruStain FcX (Biolegend), and stained with antibodies (**Table S3**). After extracellular staining, cells were washed with stain buffer (PBS, 2% FBS, 0.05% NaN3) twice prior to fixation and permeabilization using the True Nuclear Transcription Factor Buffer Set (Biolegend) according to manufacturer instructions. Cells were blocked with normal rat serum prior to staining for transcription factors overnight at 4°C in permeabilization buffer. Cells were washed with stain buffer prior to reading on a Cytek Aurora 5L cytometer. For analysis of cytokine expression, cryopreserved pLN suspensions were thawed and incubated at a concentration of 1e6/mL with Cell Stimulation Cocktail (Invitrogen) in the presence of 0.66μL/mL GolgiStop (BD) for four hours. Cells were stained with Zombie Aqua (Invitrogen), incubated with TruStain FcX (Biolegend), and stained with a panel of antibodies (**Table S4**). After extracellular staining, cells were fixed with BD cytofix and permeabilized with Intracellular Staining Perm Wash Buffer (Biolegend). Staining for cytokines was conducted overnight at 4°C in perm wash buffer. Cells were washed with perm wash buffer prior to reading on a Cytek Aurora 5L cytometer. Data were analyzed using FlowJo V10.8.2 software.

### Mass Cytometry

Cryopreserved cells from pLN of ND (n=12) and T1D donors (n=10) were stained with a 35-marker panel (**Table S5**) of MAXPAR metal-chelating polymers and run on a CyTOF mass cytometer by the Human Immune Monitoring Center at Stanford University, as reported previously^20^.

### Mass Cytometry Analysis Pipeline

Data were normalized between batches using premessa^21^ in R based on internal bead controls. After this, FCS files were loaded into FlowJo (BD Biosciences), beads were excluded, and live single CD45^+^ cells were used for downstream analysis (**Figure S1**). Data analysis involved importing FCS files into R and creating a flowset using the flowCore package^22^. Data were arcsinh transformed with a cofactor of 5 prior to using the expression data and cell metadata to create a Seurat object. Principal component analysis (PCA) was run using all panel markers, after which K nearest neighbors (KNN) were calculated using FindNeighbors(), and initial clustering using the lovain algorithm was performed (resolution=0.2). PCA was re-run on subsetted data with markers not associated with the lineage of interest removed. Nearest neighbors were calculated as above, and subclustering was performed as above for each subset using the Louvain algorithm (resolution=0.2). Cell counts per cluster per individual were determined and normalized by the total number of cells to allow for comparison of cluster proportions. Differential abundance analysis was performed using propeller^23^ with age and batch as covariates in a linear model designed with model.matrix(∼0 + group + age + Batch). Design matrix and contrasts (T1D-ND) were incorporated into the propeller.ttest function with arguments robust=T, trend=F, sort=T. Significantly altered cluster proportions were defined as Bonferroni adjusted p-value <0.05.

### Single Cell RNA Sequencing

Cryopreserved single cell suspensions were thawed and stained with a cocktail of 7 oligo-barcoded TotalSeq antibodies (Biolegend, **Table S6**) and peptide-MHC dextramers (Immudex, **Table S6**) to aid in clustering naïve and memory CD8^+^ and CD4^+^ T cell phenotypes and to aid in identification of antigen-specific cells. Prior to use, Totalseq antibodies were centrifuged at 14000x*g* for 10 minutes at 4°C to avoid antibody aggregates, as recommended by the manufacturer. Cell viability was assessed using acridine orange and propidium iodide (AO/PI) staining on a Nexcelom Cellometer or CellDrop. For six samples with viability <70% (**Table S7**), the Dead Cell Removal Kit (Stemcell) was used. 1x10^6^ cells per sample (viability >70%) were resuspended in 100μL volume of PBS + 1% BSA prior to incubation with 2μL of each pMHC dextramer according to manufacturer’s instructions for 10 minutes at room temperature (RT), followed by incubation with TruStain FcX (5μL/test) to block Fc receptors for 10 minutes at 4°C, and incubation with the antibody master mix (0.5μL of each antibody/test) for 30 minutes at 4°C. Cells were washed four times with 3mL of chilled PBS + 1% BSA, after which live cells were counted using AO/PI on a Nexcelom Cellometer or CellDrop, and volumes adjusted to obtain a targeted recovery of 10,000 cells. Single cell suspensions were loaded into the Chromium Controller (10X Genomics) and libraries prepared according to the 5’ v1.1, 5’ v2, and 5’ HT v2 kit protocols (10x Genomics). Libraries were sequenced on an Illumina Novaseq or Novaseq X instrument with a target of 50,000 paired reads for gene expression, 10,000 paired reads for surface protein, and 5,000 paired reads for TCR libraries. Median gene expression library sequencing saturation was 80.9%.

### Single Cell Read Processing

BCL files were processed to fastq files using the cellranger mkfastq pipeline (10X Genomics). Fastq files from gene expression, surface protein, and TCR libraries were processed to matrices using the cellranger multi (10X Genomics) count pipeline in order to maximize counting of cells with all three modalities.

### Single Cell RNAseq Normalization and Analysis Pipeline

scRNAseq data were subject to quality control (QC) by removing doublets using the doubletfinder package^24^, and cells were removed based on cutoffs of reads/cell ≤2000 and percent mitochondrial reads ≤10%, then normalized using the DSB package^25^ to remove background noise and ambient signal from RNA and protein libraries. After this initial filtering, data were log-normalized with NormalizeData() and scaled with ScaleData(), then variable features were calculated, and dimensionality reduction was performed in Seurat^26^. Due to a lack of observed pMHC dextramer signal, we excluded these from future analysis. For the antibody-derived tag data, variable features were set to all antibodies included, data were scaled^26^, and PCA was run. Data were integrated in Seurat using IntegrateLayers() with RPCA as the integration method. After integration of gene expression data by patient ID, we observed no significant batch effects (**Figure S2**). After subsetting T cell clusters, data were renormalized, PCA was rerun, and RPCA integration with IntegrateLayers() was run again on this subset. Integration of TCR with gene expression data was performed using scRepertoire^27^ in R. For pancreas and pLN clone comparison, clones were defined as sequences sharing an identical complementarity determining region 3 (CDR3) amino acid sequence for both TCRα and TCRβ. Differentially expressed genes (DEGs) were calculated using MAST^28^ with sex as a latent variable with a minimum expression threshold of 0.10 and reported when false discovery rate (FDR) corrected p<0.05 and log2 fold change (FC)≥0.26. Pathway enrichment analyses were performed using the top 100 most differentially expressed genes (other than for the exhausted CD8 T cell cluster for which only 53 genes met our significance thresholds) between T1D and ND donors, with FDR p<0.05 per cluster ranked by log2FC, as input into the ReactomePA^29^ package for pLN. The R package escape^30^ was used to perform single-sample Gene Set Enrichment Analysis (ssGSEA) using the integrated pancreas and pLN dataset and normalized with nFeature RNA as the scale factor. Results were reported when Benjamini-Hochberg corrected p<0.1 as noted in figure legends. IREA analysis was performed by inputting the top (p<0.05, log2FC >0.26) DEGs between T1D and ND donors per cluster using the IREA web tool (https://www.immune-dictionary.org/app/home) with cell type of “T_cell_CD8”, species of “Human“ and Method parameter of “Score”.

### Clonotype Neighbor Graph Analysis (CoNGA)

The RNA assay from the integrated pLN Seurat object was exported in h5ad format for Python. Filtered contig annotation data from each sample were merged using make_10x_clones_file_batch from the conga package in Python. The resulting merged TCR data were used along with the gene expression data as input for run_conga.py^31^ with the –all flag and batch keys set to status and nPOD identification number (ID) in order to compare the composition of clusters by clinical status and across all donors. Gene expression data were also batch corrected using the argument batch_integration_method harmony with batch_integration_key batch. For CONGA analyses, the cellranger VDJ clonotype definition of identical CDR3 nucleotide sequence was used. Clones that possessed a conga score<1, indicative of significant overlap between gene expression and TCR neighborhoods beyond what would be expected by chance^31^, were used as input to evaluate enrichment in cluster composition by clinical status using two-sided Fisher’s exact test as reported in the figure legend.

### Phenocycler multiplex imaging

Immunofluorescence of pancreas with embedded lymph node was performed using the PhenoCycler-Fusion System (Akoya Biosciences/Quanterix) according to the vendor’s recommended protocol for formalin fixed paraffin embedded (FFPE) tissue. Deparaffinization was performed by baking at 60°C followed by xylene washes. Tissue rehydration was accomplished by moving the slide through graded ethanol/DI-H_2_O washes (100%x2, 90%, 70%, 50%, 30%, DI-H_2_Ox2). Antigen retrieval was performed by immersing sections in 1X antigen retrieval buffer under high pressure in a pressure cooker. Tissue was then washed in Hydration Buffer and then Staining Buffer. Primary antibody staining was performed using 175µL of antibodies (**Table S7**), blocking reagents (N, G-v3, J, S), and blocking buffer cocktail at RT. The tissue was then rinsed in Staining Buffer and sequentially post-fixed, followed by PBS washes, in 1.6% paraformaldehyde, then ice-cold methanol, and finally Fixative Solution. Tissue was then kept overnight in Storage Buffer at 4°C. The following day, a 96-well reporter plate was prepared with either cycle-specific cocktails of complementary oligonucleotide-conjugated fluorescent secondaries, Nuclear Stain (DAPI), and Reporter Stock Solution for cyclic protein detection or only Reporter Stock Solution for blank cycle autofluorescence detection. Finally, a Flow Cell was press-sealed to the face of the slide for microfluidic delivery of reagents.

Automated detection and imaging were performed by first generating an experiment file specifying target-barcode pairs and exposure times for each cycle (Experiment Designer software, Akoya Biosciences/Quanterix). Phenocycler Buffer and dimethyl sulfoxide solutions (20% and 90% in 1X Phenocycler Buffer) were prepared and loaded in the Phenocycler-Fusion instrument bottles. Following automated calibration and equipment checks, tissue was auto-detected, and images were acquired with 16-bit precision at 20x magnification, automatically processed for stitching and autofluorescence subtraction, and saved in the proprietary ‘qptiff’ format.

### Phenocycler data analysis

Semantic segmentation was performed in QuPath^32^v6.0 T on the qptiff file for donor 6551. Objects (pLN/Pancreas tissue area and islets) were classified using the pixel classifier with default settings. Cell and nuclear boundaries were generated with the Instanseg^33^v0.1.5 extension’s default model “fluorescence_nuclei_and_cells-0.1.1” using all channels as input, default cell measurements, and default outputs. Statistics for each cell’s shape and marker intensity, as well as centroid coordinates and signed distance to nearest islet, were exported as a csv file for analysis in a custom Python Jupyter notebook^34^.

### Statistics

Data were analyzed using R v4.2.3, python v3.9 and GraphPad Prism v9. Statistical tests and p-values are reported in the figure legends.

### Study approval

The study was conducted in accordance with federal guidelines and the University of Florida (UF) Institutional Review Board approved protocol IRB201600029, as previously described^35^.

### Data and code availability

Mass cytometry and flow cytometry data are available at ImmPort (https://www.immport.org, accession number: SDY3066). Supporting Data Values associated with this manuscript are provided in the supplement. Gene expression and TCR data are available on GEO with accession: GSE298811. Phenocycler data are available upon request, and code used for Phenocycler analysis can be accessed at https://github.com/smith6jt-cop/Panc_pLN_Analysis.

## Results

### Mass cytometry reveals a population of CD8^+^CXCR3^+^ T cells enriched in T1D pLN

The diverse nature of immune cell phenotypes mandates the use of high-parameter technologies, such as cytometry time of flight (CyTOF)^36, 37^ ^12^, to deep-phenotype multiple immune subsets. We applied high-parameter CyTOF to pLN derived from 10 T1D and 12 ND nPOD organ donors (**Table S8, Figure 1A**). Clustering total CD45^+^ immune cells revealed 11 unique populations spanning B cells, CD4^+^ T cells, CD8^+^ T cells, myeloid, and natural killer (NK) cells (**Figure S3A**). We observed no significant differences in cluster abundance globally between T1D and ND donors (**Figure S3B**);

**Figure 1.**
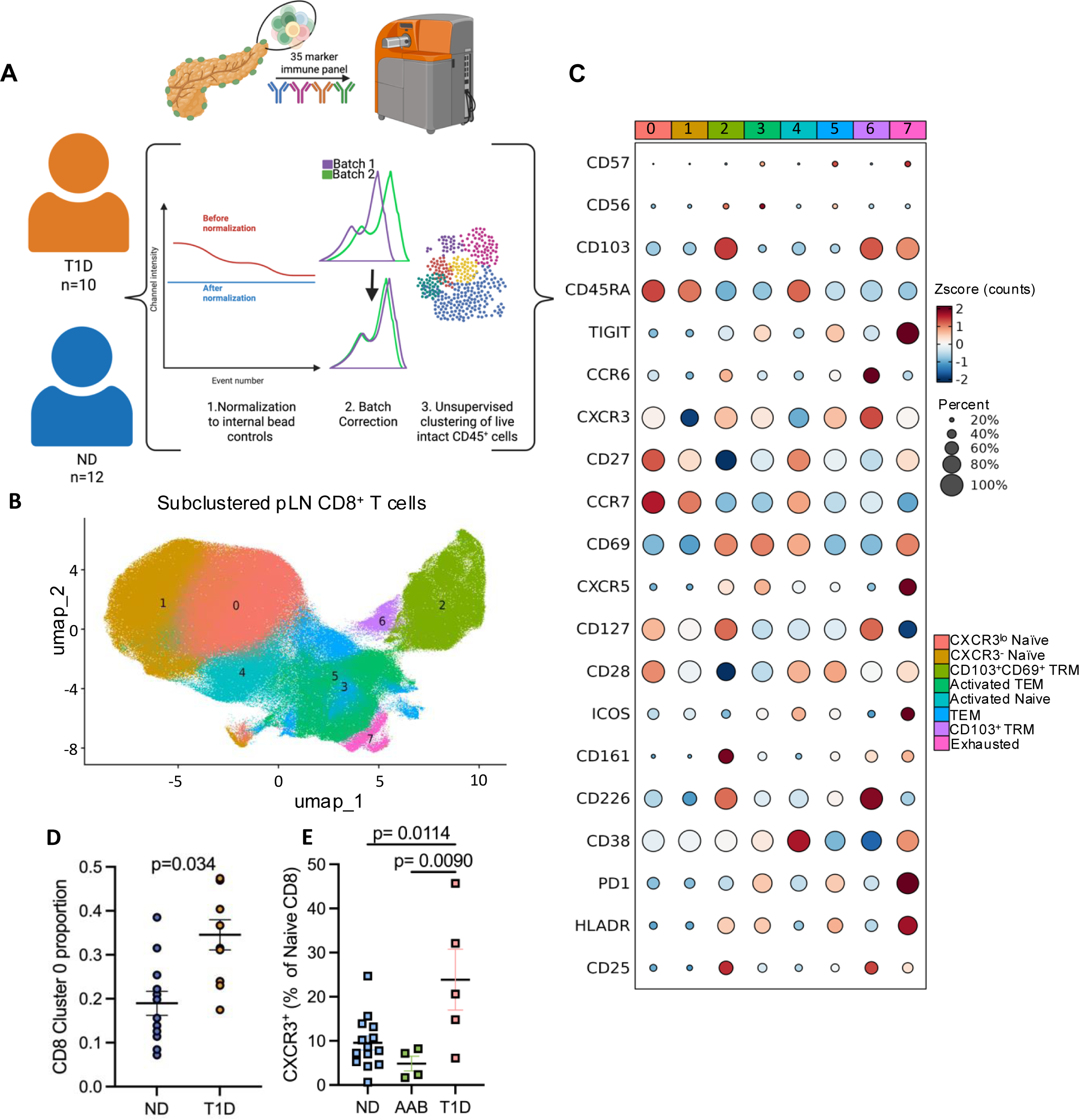
Differential abundance of CXCR3^lo^ naïve CD8 T cells in T1D pLN. A) Experimental workflow involving processing of pancreatic lymph node (pLN), staining for lineage, phenotypic, and quality control (QC) markers, and data collection on a mass cytometer. After data acquisition, normalization of channels to internal bead controls was performed using the premessa^21^ package, and any confounding effects of batch or donor age on cluster composition were accounted for in analysis through incorporation as covariates in a linear model. B) UMAP of subclustered CD8^+^ T cells. C) Z-scored DotPlot of marker expression for each CD8^+^ T cell subcluster. D) Significantly increased proportion of subcluster 0 (CXCR3^lo^ naïve like) in T1D pLN as determined by linear modeling with donor age and CyTOF batch as covariates. E) Significantly increased percentage of naïve CD8^+^ T cells expressing CXCR3 found in an independent cohort of pLN donors by flow cytometry. P value for (E) is the result of an ordinary one-way ANOVA with Tukey’s multiple test correction.

We performed subclustering to obtain a finer resolution of immune cell subsets from the CyTOF data. As one batch demonstrated slight, non-significant differences in age (**Figure S4**), we chose to examine differences in cluster proportions using a linear model with age and batch incorporated as covariates. Subsetting and reclustering of B cells and CD4 T cells yielded no significant changes in cluster abundance or marker expression by clinical status (**Figure S5**). However, CD8^+^ T cell subclustering yielded eight proteomic clusters (**Figure 1B**), with a T1D-associated increase in the proportion of CD8^+^ T cell cluster 0, expressing the naïve T cell markers CD45RA and CCR7, costimulatory molecules CD27 and CD28, as well as low levels of the chemokine receptor CXCR3, which confers migratory capacity towards CXCL10^38^ (**Figure 1C-D**). Though we observed more variability due to the small number of donors, this finding was also confirmed in an independent flow cytometry cohort consisting of 23 nPOD pLN cell isolates from ND, autoantibody positive (AAB+) and T1D donors (**Table S9**) (**Figure 1E, Figure S6**), where T1D donors possessed a significantly increased proportion of CXCR3^+^ cells within the naïve (CD45RA^+^CCR7^+^) CD8^+^ T cell compartment as compared to both ND (p=0.0114, FC=2.50) and AAB+ donors (p=0.0090, FC=4.91). Importantly, we did not observe an association of the frequency of this cluster or the related flow cytometry phenotype with age or disease duration (**Figure S7A-D**). These phenotypic signatures are clinically relevant in peripheral blood studies: we recently reported a T1D-associated increase in CXCR3^lo^ naïve CD8^+^ T cells^19^, and circulating CD8^+^ T cells with a more terminally differentiated/exhausted profile have been previously correlated with preservation of C-peptide and response to teplizumab^12, 13^.

### IL-15 drives memory maintenance and averts the exhaustion program in T1D CD8^+^ T cells

To explore the transcriptomic profile of the cell populations previously identified by both CyTOF and fluorochrome-based flow cytometry, we performed single cell RNA sequencing (scRNAseq) analysis with TCR-seq on donor pLN (**Figure 2A**). After QC, we obtained 122,300 cells across 16 individuals (9 T1D, 7 ND) (**Table S10**) that comprised 38 clusters spanning CD4^+^ T cells, CD8^+^ T cells, B cells, NK cells, innate lymphoid cells (ILCs), monocyte/macrophages, and dendritic cells (DCs) (**Figure 2B**, **Figure S8A-B**). No significant differences in global cluster abundance across clinical status were detected (**Figure S8C-D**). We performed subclustering to more deeply examine alterations in T cell phenotype, yielding 19 subclusters spanning naïve, memory, and exhausted T cells, as well as Tregs (**Figure 2C**). Differential expression analysis of T1D compared to controls identified conserved signatures across multiple CD4 T cell clusters, such as a reduction in genes responsible for negative regulation of cytokine signaling (e.g., *TNFAIP3*^39^ within naïve CD4, T follicular helper [TFH], Th1/17, thymic Treg [tTreg] clusters) and activation/proliferation (e.g., *JUND*^40^ within TFH, tTreg, Th1/17, Th17 clusters), as well as a broad increase in interferon signaling genes (e.g., *IFITM3*^41^ within Th17, naïve CD4, exhausted CD4, activated naïve CD4, TFH, Th1/17) (**Table S11**).

**Figure 2.**
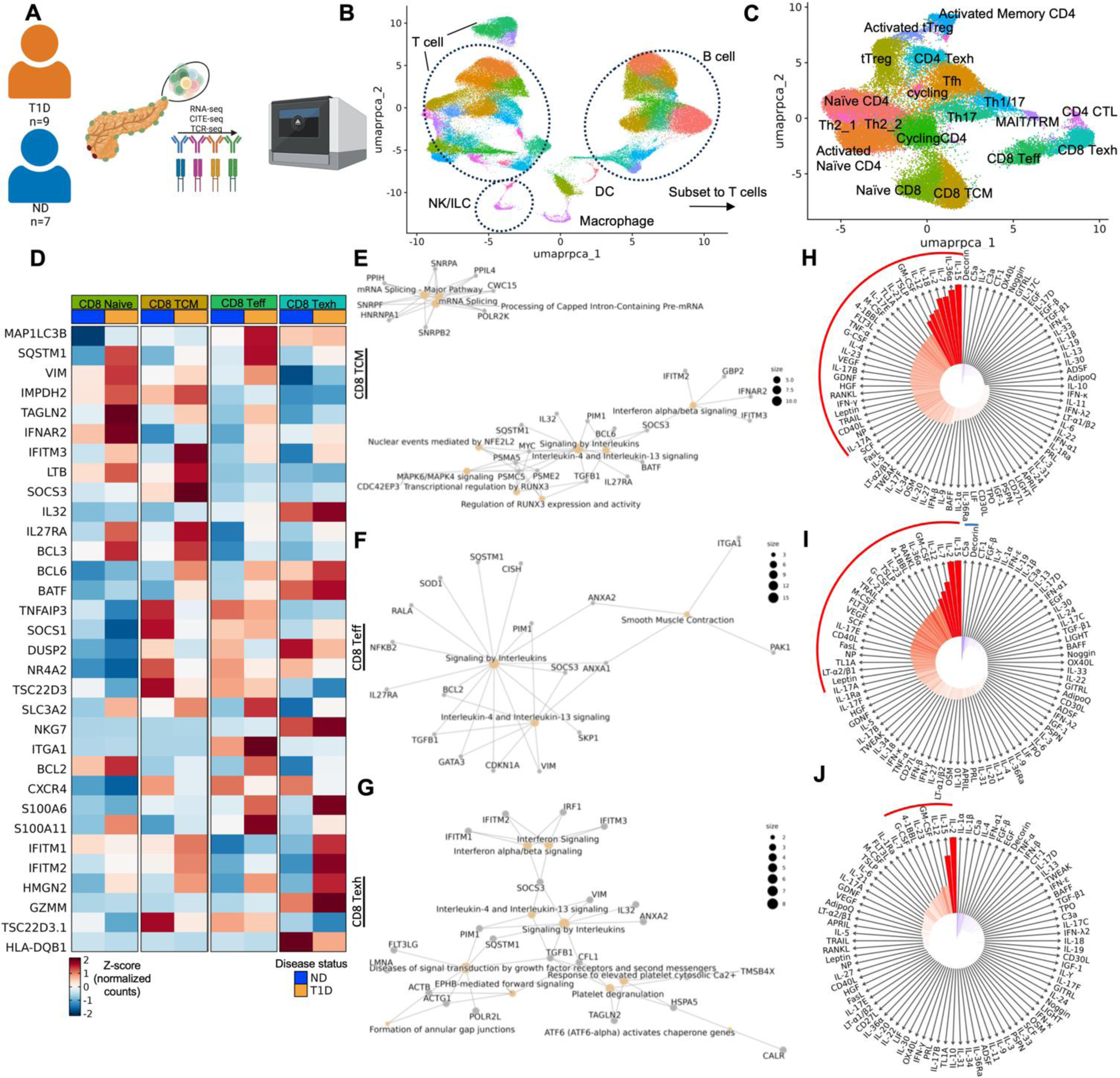
IL-15 is predicted to drive signatures of memory maintenance and effector function in T1D. A) Experimental workflow for scRNAseq of pLN. B) UMAP projection of total pLN cells, which were subsetted to T cells, reintegrated with rpca, and reclustered, resulting in 19 T cell clusters (C). D) Heatmap of Z-scored gene expression data showing a selection of DEGs between T1D donors and ND donors across naïve, Teff, and Texh CD8^+^ T cells. E) Network plot using DEGs in naïve CD8 T cells showing enrichment of reactome pathways in T1D. F) Network plot using DEGs in CD8 Teff cells showing enrichment of reactome pathways in T1D. G) Network plot using DEGs in CD8 Texh cells showing enrichment of reactome pathways in T1D. H) Compass plot of IREA results using the top genes upregulated in T1D in naïve CD8^+^ T cells (p<0.05, log2FC>0.26) as input. I) Compass plot of IREA results using the top genes upregulated in T1D in CD8^+^ Teff cells (p<0.05, log2FC>0.26) as input. J) Compass plot of IREA results using the top genes upregulated in T1D in CD8^+^ Texh cells (p<0.05, log2FC>0.26) as input. H-J) Bar height is indicative of enrichment score, while color corresponds to FDR-adjusted P value (two-sided Wilcoxon rank-sum test), red or blue arcs indicate significant positive or negative enrichment, respectively. Genes highlighted on heatmap (D) met log2FC cutoffs of 0.26 and adjusted p-value cutoffs of 0.05. Pathway analysis was performed with the reactomePA package and visualized if adjusted p<0.1 with the clusterprofiler package.

We next focused more deeply on CD8^+^ T cell subsets as these phenotypes were significantly altered in our CyTOF and flow cytometry data. When comparing DEGs (defined as FDR corrected p<0.05 and Log2FC≥0.26) between T1D and ND pLN, we observed within the naïve CD8^+^ T cell cluster (**Figure 2D**) an upregulation of genes involved in autophagy (*MAP1LC3B*^42^, *SQSTM1*^43^), migration (*VIM*), proliferation (*IMPDH2*^44^), and cytokine production (*TAGLN2*^45^). CD8 TCM cells from T1D donors showed upregulation of inflammatory cytokine signaling (*IFNAR2*^46^, *IFITM3*, *LTB*^47^, *SOCS3*^48^, *IL32*^49^, *IL27RA*^50^) as well as regulators of terminal differentiation (*BCL3*^51^, *BCL6*^52^, *BATF*^53^). Moreover, both naïve and TCM CD8^+^ T cells from T1D donors also downregulated genes involved in negative regulation of cytokine signaling (*TNFAIP3*^39^, *SOCS1*^54^) and genes associated with T cell exhaustion (*DUSP2*^55^, *NR4A2*^56^, *TSC22D3*^57^) as compared with ND. In the CD8 effector T cell (Teff) cluster (**Figure 2D**), we observed an upregulation of genes involved in effector function (*SLC3A2*^58^*, NKG7*^59^), as well as migratory genes (*VIM*^60^*, ITGA1*^61^) in T1D relative to ND donors. We also noted upregulation of the anti-apoptotic factor *BCL2*, which is important for the maintenance of long-lived memory subsets^62, 63^, within CD8 Teff in T1D. Moreover, within this cluster, we observed downregulation of genes associated with terminal differentiation and exhaustion (*CXCR4*^64^, *DUSP2*^55^), indicative of restraining of terminal differentiation in T1D as compared to controls. Similarly, in the exhausted CD8 T cell cluster (CD8 Texh) (**Figure 2D**), we observed upregulation of genes involved in migration (*VIM*^60^), calcium signaling (*S100A6*, *S100A11*^65^), inflammatory cytokine response (*SOCS3*, *IFITM1*, *IFITM3*, *IFITM2*^41, 66^), and cytotoxicity (*HMGN2*^67^, GZMM), while genes associated with terminal differentiation and exhaustion (*TSC22D3, DUSP2*, *HLA-DQB1*^68^) were downregulated in T1D. We did not detect differential expression of other canonical markers of stemness or exhaustion (i.e., *TCF7*, *LEF1*, *CD122*, *CXCR3*, *PD1*), potentially due to data sparsity or differential regulation of protein and RNA expression.

Pathway analysis using the Reactome database revealed that naïve CD8 T cells (**Figure S9A**) in T1D were enriched for autophagy and mitophagy. Additionally, CD8 TCM cells in T1D were enriched for interleukin signaling and interferon signaling (**Figure 2E**), while CD8 Teff (**Figure 2F**) were enriched for interleukin signaling and smooth muscle contraction, the latter likely reflective of increased calcium signaling (*ANXA1*, *ANXA2*)^69^. Lastly, CD8 Texh (**Figure 2G**) were enriched for interferon signaling, interleukin signaling, and signal transduction through growth factor receptors (*TGFB1*, *FLT3LG*^70^) and second messengers (*PIM1*^71^). We then performed Immune Response Enrichment Analysis (IREA), which uses an extensive database of specific gene sets induced by cytokine treatment^72^, to infer the cytokine milieu driving these CD8 phenotypes. Across the trajectory of CD8^+^ T cell differentiation, IL-15 signaling was notably enriched in T1D (**Figure S9B**; **Figure 2H-J**), a pathway that is known to promote stemness and restrain exhaustion-like phenotypes^73^ and can synergize with other inflammatory cytokines to promote effector function^74^. Collectively, these data and our prior peripheral blood findings^75^ suggest Tc1 skewing as an immunological feature of T1D shared across tissues.

### CD8^+^ T cell clones possessing naïve-like and *TCF7*^+^*TOX*^+^ phenotypes are more prevalent in T1D pLN

Separate analyses of receptor or transcriptomic data may preclude insights into the relationships between repertoire characteristics and phenotype. Thus, recent methods have been developed to enable joint analysis of these modalities, namely Clonotype Neighbor Graph Analysis (CoNGA)^31^, which has shown utility in revealing transcripts associated with specific TCR genes and features. We applied CoNGA to our pLN dataset, which identified 35 groups with significant (CoNGA score <1) overlap in TCR and gene expression neighborhoods and a minimum of 4 cells per group (**Figure S10**). In accordance with the original CoNGA publication^31^, we found a stronger relationship between receptor and phenotypes in CD8^+^ T cells and invariant T cells, with the majorityof our CoNGA hit clusters being comprised of CD8^+^ T cells, including naïve, effector, and mucosal-associated invariant T (MAIT) cell clusters (**Figure S10**). Among these clusters, we found 10 that were overrepresented to some degree in T1D donors (**Figure 3A**, indicated by leftmost orange bar). In particular, we observed that CoNGA clone hits belonging to CoNGA gene expression (GEX) cluster 8 and TCR cluster 4 were found in several T1D donors, the highest frequency being found in new-onset donor 6550, while 37/38 clones detected in ND donors (97%) were found in a single ND donor possessing high risk HLA alleles (HLA A*0201, HLA DRB1*0401) (**Table S12)**. Thus, clones belonging to these clusters were more often found in T1D (p=0.0012, OR=1.988) (**Figure 3B**). The enriched GEX cluster 8/TCR cluster 4 is comprised of CD8^+^ T cells with high TCR diversity (113 unique clonotypes and 113 total cells, indicative of no clonal expansion) and with high expression of *CCR7*, *SELL*, *LEF1*, *TCF7*, and *IL7R*. We also noted that this cluster possessed some features suggesting prior activation, namely *KLRK1*^76^, costimulatory molecule *HCST*^77^ encoding DAP10, and low *CXCR3* expression. The TCRs within this cluster possessed a diverse TCR beta chain (*TRB*) repertoire, while alpha chain (*TRA*) gene usage was restricted to TRAV14/DV4, TRAV38-2/DV8, and TRAV38-1 (**Figure 3A**). Additionally, this cluster also possessed a low Atchley factor 4 score (blue af4 logo, TCRseq features panel, **Figure 3A**), which has previously been associated with larger, more hydrophobic residues^31^. In addition to this naïve-like cluster, we also noted enrichment of a less frequent cluster (GEX cluster 11, TCR cluster 13), which possessed a more terminally differentiated profile, expressing cytotoxicity markers (*GZMA*, *GZMK*, *NKG7*), costimulatory molecule *HCST*, as well as *CXCR3* and both *TCF7* and *TOX* (**Figure 3A**). Though infrequent, this cluster was also present in multiple T1D donors, with only 2/18 clones being found in ND donors. We searched for known T1D clones^78^ in these T1D-associated CoNGA clusters based on similarity to known alpha chain complementary determining region 3 (CDR3) sequence, given recent data on potential alpha chain biases in autoreactive clones in T1D^79^. We found within the naïve-like cluster several clones which share CDR3a (allowing for one amino acid mismatch) with known T1D clones, namely four GAD65 reactive clones, three preproinsulin reactive clones, and a clone which possessed a CDR3a chain shared across multiple specificities in a new onset T1D donor. We also found three clones with matching CDR3a to clones with GAD65, ZNT8_186-194_ and PPI_16-25_ specificities in a control donor with high-risk HLA alleles (**Table S12**). In the more terminally differentiated cluster, we found an expanded clone (clone size of 6) matching a PPI reactive clone, as well as sharing an alpha chain motif “AGGYQKVTF” present in the previously published 22.A10^80^ PPI_1-11_ reactive clone (**Table S12**). Thus, these enriched clusters may contain autoreactive specificities. Interestingly, we also noted an increase (p=0.0007, OR=2.153) in a group of expanded TCRs (CoNGA gene expression cluster 16 paired with TCR cluster 13) comprised of MAIT and semi-invariant T cells (*TRAV1-2* gene usage paired with a more diverse set of *TRB* chains) in T1D pLN (**Figure S11**).

**Figure 3.**
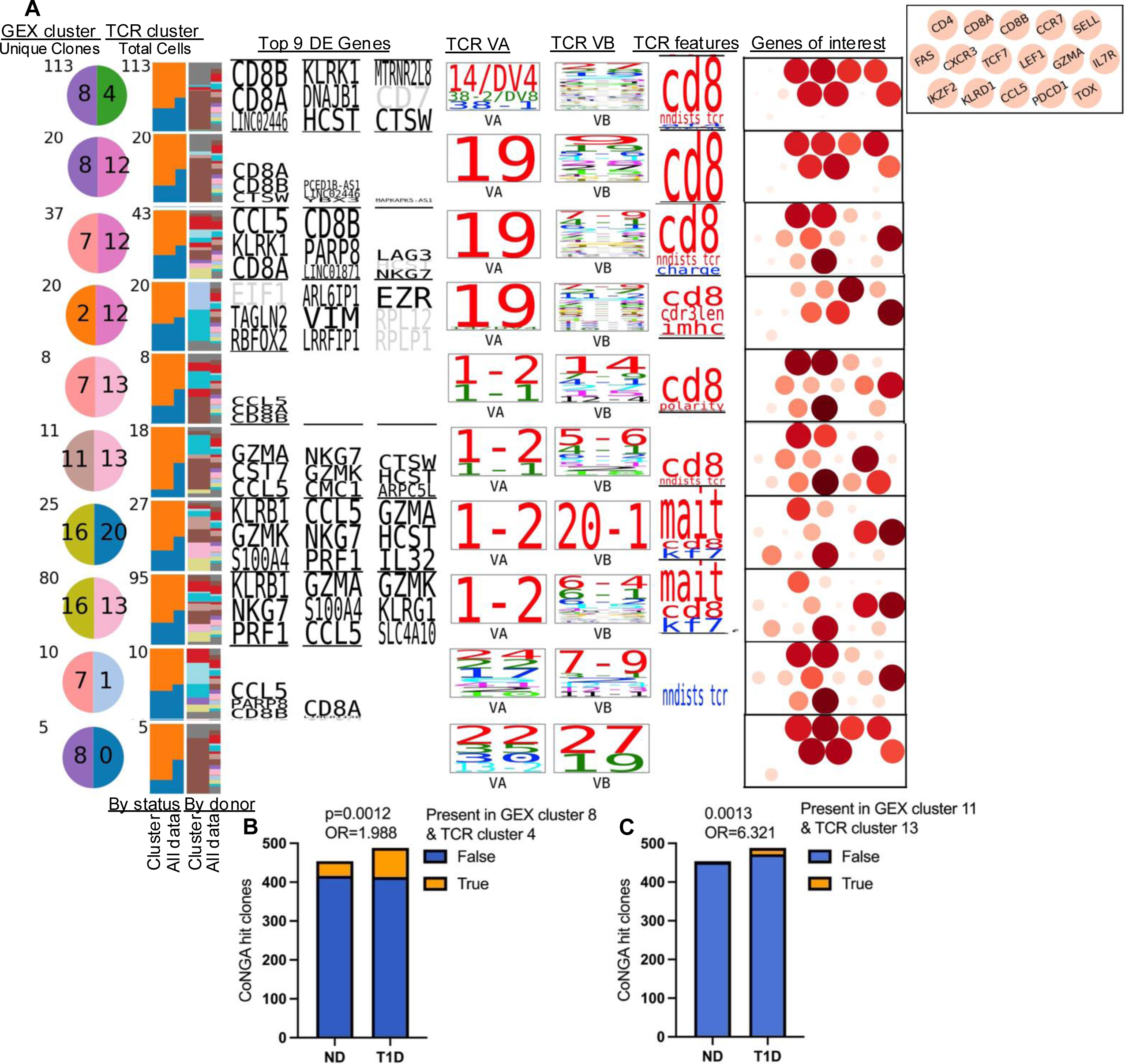
Integrative receptor and gene expression analysis highlights an enriched naïve CD8^+^ T cell TCR cluster in T1D pLN. A) Gene expression and TCR Graph vs Graph results for CoNGA clusters with score <1. Left half of semicircle indicates gene expression cluster, right half of semicircle indicates TCR cluster assignment. Bar plots indicate the relative proportion of T1D (orange) and ND (blue) cells for the given cluster (left) and for the entire dataset (right), followed by the same for nPOD ID, where each color represents a different donor. Top 9 differentially expressed genes are shown per cluster, as well as TCR genes, and TCR sequence features^31^ (TCRseq features panel, with red indicating higher and blue indicating lower scores relative to the rest of the dataset). A panel of manually curated marker genes shown is shown on the far right with red dots colored by mean expression and sized by percent of cells expressing the gene. DEGs are determined by Wilcoxon test and scaled by adjusted p value, where full height requires p<10^-6^. Fold changes <2 are shown in grey. Difference in cluster proportion assessed with two-way Fisher’s exact test.

### Shared memory T cell populations demonstrate enhanced effector phenotype in pancreatic slices

To investigate receptor sharing and gene expression across tissues, we performed scRNAseq on isolated immune cells from fresh pancreatic slices^81, 82^ and paired pLN (**Figure 4A**) from 2 recent-onset T1D donors (nPOD 6551, duration 0.58 years; nPOD 6536, duration 4 years) with independently confirmed insulitis^83^. Data integration yielded 17 clusters, which included naïve, memory, and effector CD4^+^ and CD8^+^ T cell populations, Tregs, B cells, and resident macrophage populations (**Figure 4B**). We first interrogated the TCR repertoire to examine sharing of T cell clones across these tissues, and while both donors possessed insulitis, the newer onset donor 6551 possessed more extensive insulitis than donor 6536 (**Figure 4C**). We found that expanded clones localized mostly to clusters 0 (CD8 Teff) and 2 (CD8 CTL), respectively corresponding to CD8^+^ T cell populations with lower and higher expression of cytotoxicity-related molecules (*GZMK*, *CST7*, *LTB*) (**Figure 4D-E**). Clone sharing across pancreas and pLN was minimal, with morisita horn (MH) values of 0.007 and 0.002 for cross-tissue overlap for donors 6536 and 6551, respectively (**Figure 5F**). However, even with our limited sample size, we were able to detect 6 unique clonotypes shared across tissues and clusters, with 2 clones shared across tissues (**Figure 4G**, **Table 1**). We detected an expanded CD4^+^ T cell clone in donor 6536, which was shared between tissues (cluster 9, ICOS^hi^ TCM), as well as an expanded CD8^+^ T cell clone in donor 6551, which was shared between tissues (clusters 0/2 CD8 Teff and CTL). The majority of sharing was detected between the effector CD8 clusters 0 and 2, especially within donor 6551, as compared to donor 6536, perhaps due to reduced infiltration as a result of increased disease duration.

**Figure 4.**
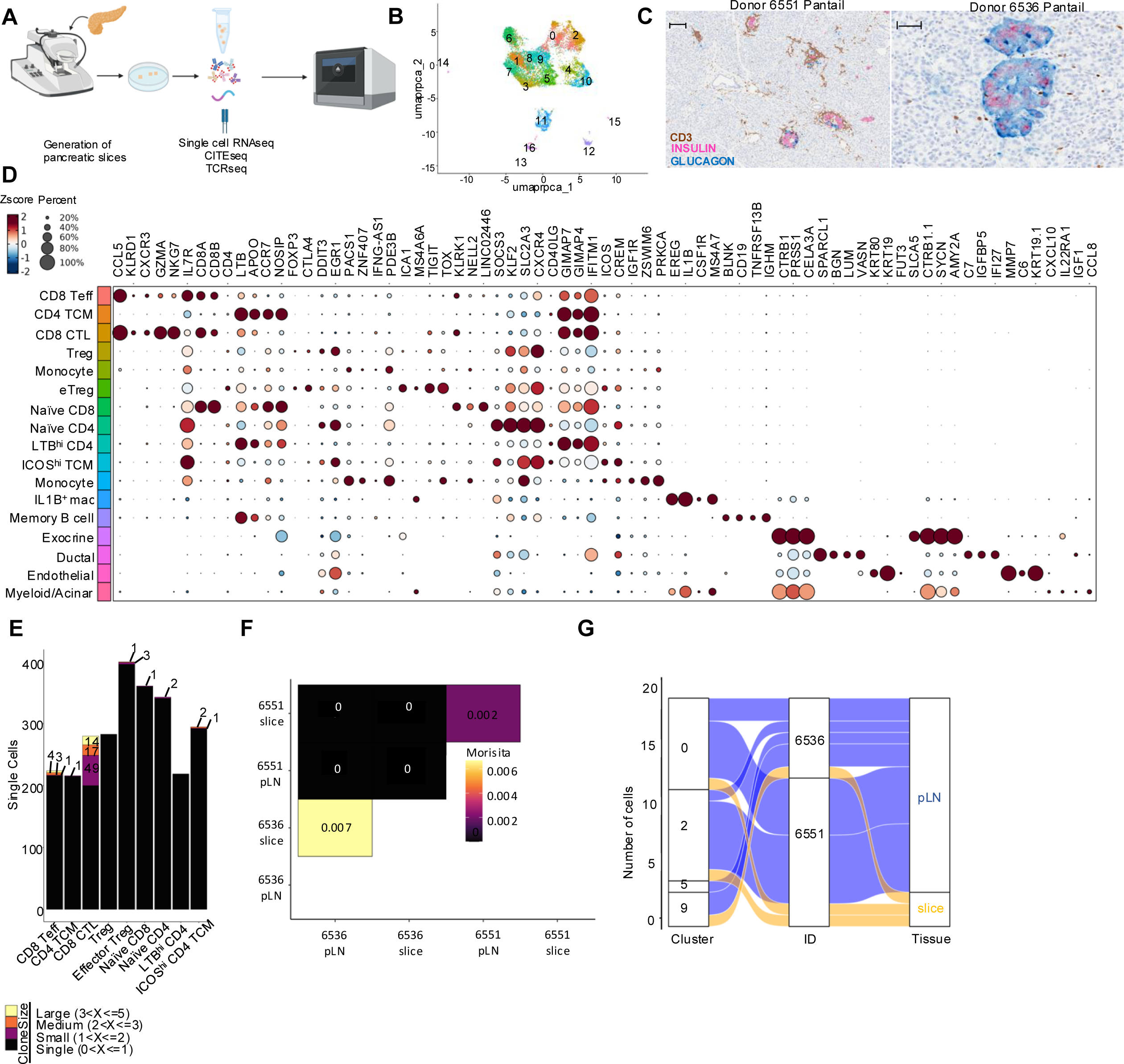
Cross-tissue clone and phenotype sharing in human pLN and pancreas. A) Processing workflow of pancreatic slices where pancreas was embedded in agarose and sliced using a vibratome to yield 150μm slices^81, 82^, after which cells were processed to single cell suspension, selected for CD45^+^ cells, and run on the 10X Chromium controller as described in methods. B) UMAP of integrated gene expression data (n=2 donors, pLN and pancreatic slices). C) Histological staining (obtained from nPOD Aperio e-slide manager) of insulin (pink), glucagon (blue), and CD3 (brown) across pancreas tail of the two donors with paired pancreatic slice and pLN data shows increased insulitis in new-onset donor 6551. Scale bars are included in the upper left corner of each image and correspond to 10x magnification and 101um for donor 6551, and 20x magnification and 50um for donor 6536. D) Dotplot showing Z-scored gene expression per cluster in (B), with dot size corresponding to percent of cells in the cluster expressing the marker. E) Barplot illustrating distribution of expanded clones across clusters, with the majority of expanded clones localizing to memory CD8^+^ T cell clusters 0 (CD8 Teff) and 2 (CD8 cytotoxic T lymphocytes [CTL]). F) Assessment of repertoire overlap across tissues with the Morisita-Horn metric shows limited sharing across tissues. G) Alluvial plot subsetted to only shared clones showing clonal sharing between clusters 5 (eTreg) and 9 (ICOS^hi^ TCM), and 0 (CD8 Teff) and 2 (CD8 CTL) across both clusters and tissues.

**Figure 5.**
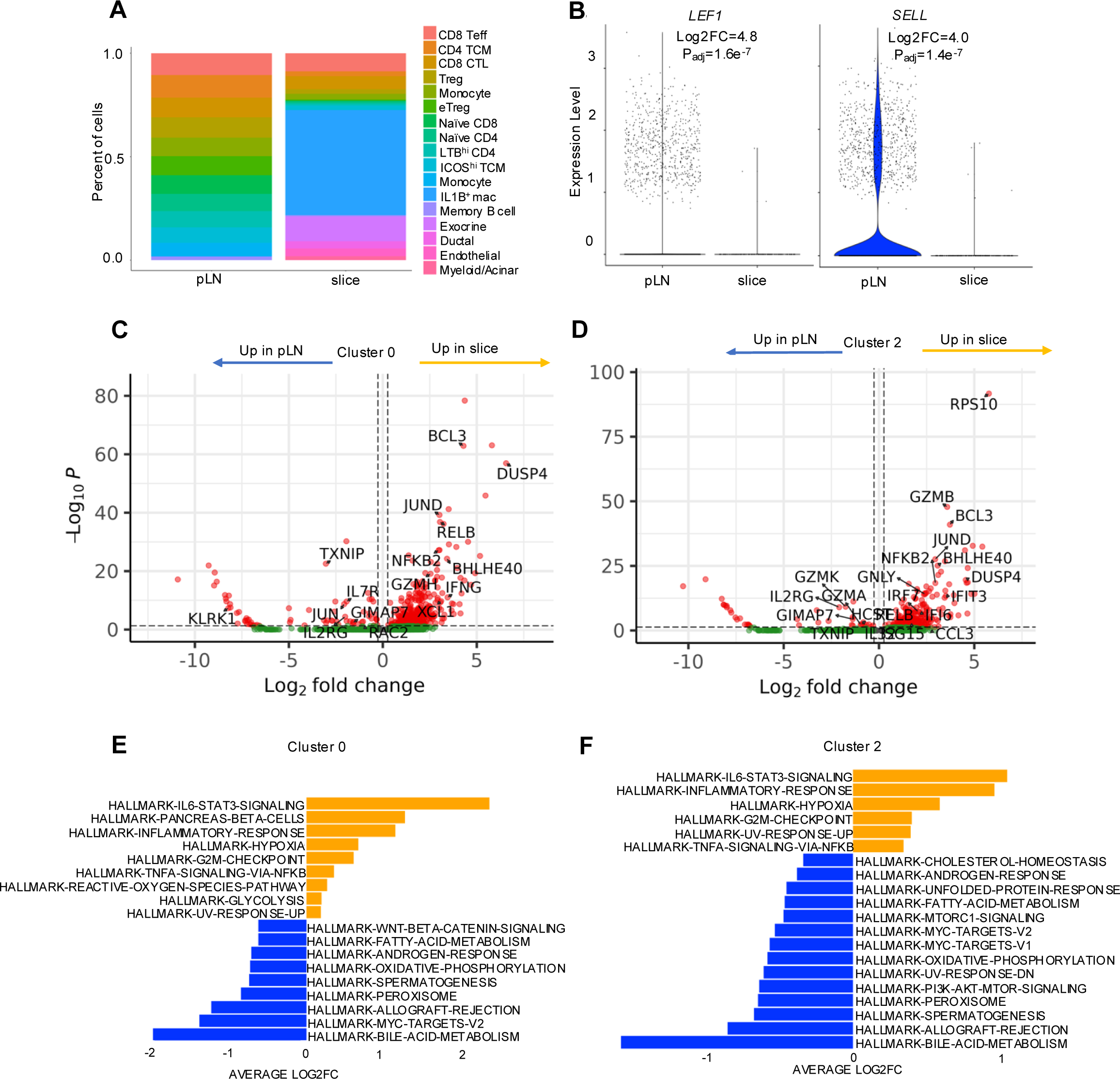
Tissue-dependent gene expression programs of memory CD8^+^ T cell populations. A) Barplot of percent of each cluster per tissue illustrates differential subset abundance of innate and adaptive immune cells. B) Violin plots illustrating differential expression of *LEF1* and *SELL* in CD8 T cells between tissues. C) Volcano plot of differentially expressed genes in cluster 0 (CD8 Teff) between pancreatic slices and pLN. D) Volcano plot of differentially expressed genes in cluster 2 (CD8 CTL) between pancreatic slices and pLN. DEGs were defined as log2FC>0.26 and corrected p value<0.05. E) GSEA analysis of cluster 0 using Hallmark genesets. F) GSEA analysis of cluster 2 using Hallmark genesets. GSEA analysis was performed using escape, and DE genesets were identified with a Wilcoxon test. The top 20 gene sets with adjusted p <0.05 were plotted with positive fold change indicating upregulation in pancreatic slice (orange) and negative fold change indicating upregulation in pLN (blue).

**Table 1.**
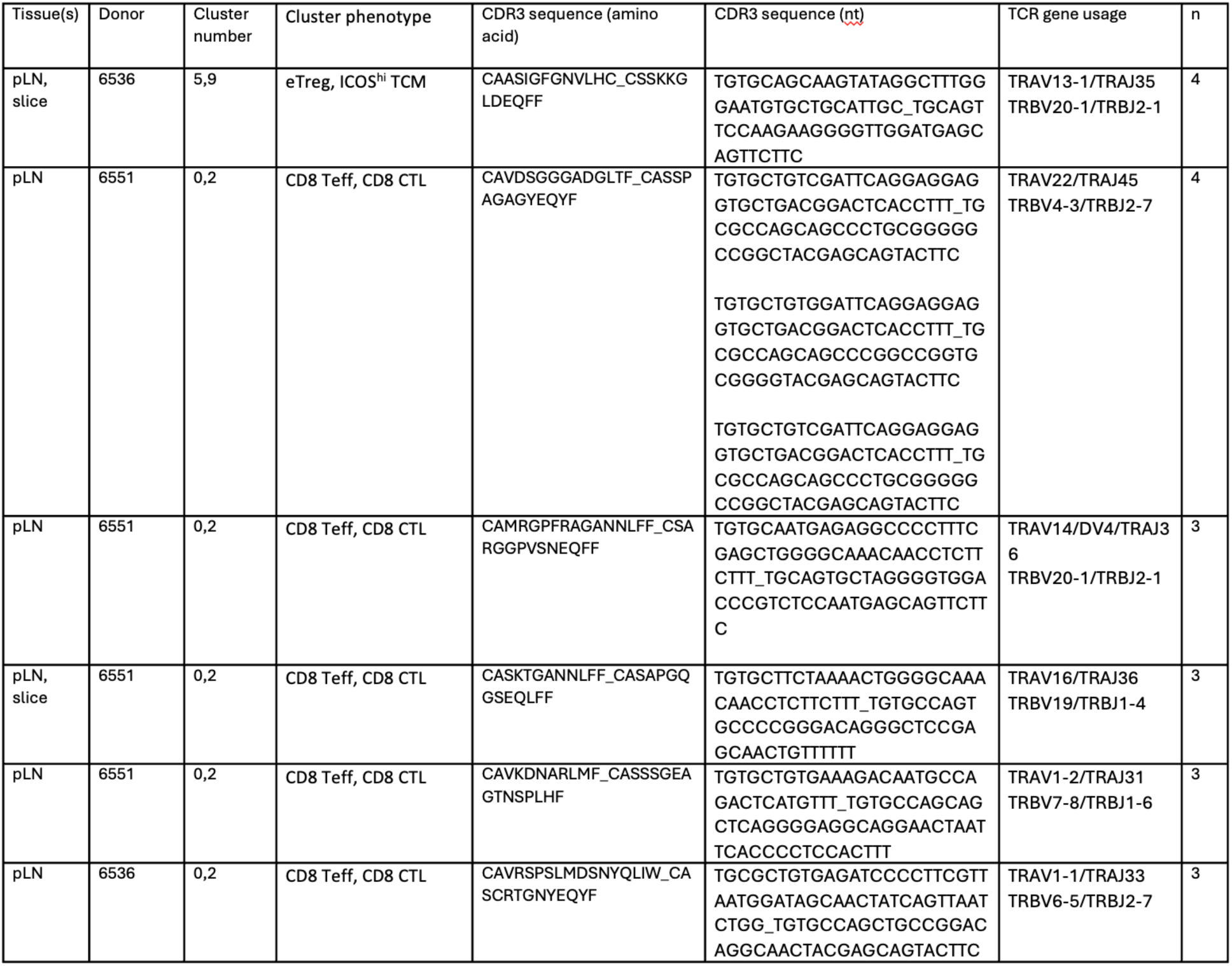
Sequences of shared clones across pLN and pancreas slices.

We next investigated tissue-specific phenotypic differences in these clusters. We noted that CD8^+^ Teff clusters 0 and 2, defined by expression of NK cell receptors (*KLRK1*^84^*, KLRB1*^85^), Th1 related chemokine receptors and ligands (*CXCR3*, *CXCR6*, *CCL5*^86^), and markers of terminally differentiated memory subsets (*LAG3*, *KLRD1*, *EOMES*^87^), were the most commonly shared clusters across tissues, while less differentiated subsets, such as naïve T cells, were expectedly more prevalent in pLN. Additionally, inflammatory macrophages expressing *IL1B* were more prevalent in pancreas (**Figure 5A**). Examining DEGs of total CD8 T cells between tissues revealed increased expression of genes involved in promoting naïve/stem-like phenotypes, *LEF1* (Log2FC=4.8, p_adj_=1.6e^-7^) and *SELL* (Log2FC=4.0, p_adj_=1.4e^-^7), in pLN relative to pancreas (**Figure 5B**). As this could partially be explained by the increased proportion of naïve cells in pLN relative to pancreatic slice, we sought to compare gene expression of shared populations across tissues. We found that cells from cluster 0 (CD8 Teff) derived from pancreatic slices expressed enhanced markers of cytotoxicity, such as *GZMB* (Log2FC=5.46, p_adj_=1.4e^-46^) as well as exhaustion marker *DUSP4*^88^ (Log2FC=6.6, p_adj_=1.3e^-57^) alongside reduced expression of common gamma chain cytokine receptors *IL7R (*Log2FC=-1.9, p_adj_=2.5e^-10^) and *IL2RG* (Log2FC=-2.0, p_adj_=4.1e^-5^) (**Figure 5C**). Cluster 2, which possessed a similar transcriptomic profile to cluster 0 but with enhanced expression of cytotoxicity and memory markers (CD8 CTL), demonstrated similar transcriptomic expression differences as cluster 0 across tissues (**Figure 5D**). GSEA of cluster 0 revealed upregulation of inflammatory signaling pathways, namely IL-6-JAK-STAT3 (Log2FC=2.3, p_adj_=1.5e^-19^) and TNFα signaling (Log2FC=0.34, p_adj_=1.7e^-7^), as well as downregulation of Wnt-Beta catenin signaling (Log2FC=-0.6, p_adj_=2.3e^-3^) (**Figure 5E**) in pancreatic slices compared to pLN, altogether suggesting that this cluster acquired a more differentiated transcriptomic program in pancreas relative to pLN. We observed similar upregulation of inflammatory pathways in cluster 2, representing an activated effector phenotype (**Figure 5F**). Notably, while Wnt-Beta catenin signaling was not significantly enhanced in pLN for this cluster, we did observe upregulation of *GIMAP5*^89^ (Log2FC=-3.3, p_adj_=1.5e^-8^) in pLN, which is an important negative regulator of GSK3β and thus, potentially enhances Wnt-signaling. In line with prior observations of IL-15 signaling in the pancreas, IREA analysis of DEGs from CD8^+^ T cell clusters 0 and 2 between pLN and pancreas revealed enriched IL-15 signaling as a driver of differential tissue phenotypes (**Figure S12**).

### CXCR3^+^ naïve T cells show features of stemness and polyfunctionality in the pLN

In order to more deeply understand the phenotype and function of the naïve CXCR3^+^ cells (cluster 0) we identified as being enriched in T1D pLN, we examined the expression of canonical exhaustion and TSCM markers on pLN CD8 T^+^ cells in a cohort of 5 T1D and 5 control donors by flow cytometry. We confirmed that CXCR3^+^ naïve CD8^+^ T cells, along with the remainder of the naïve cells (CXCR3- naïve), express significantly more TCF1 (p=0.0003, FC=2.04), and possess a significantly greater fraction of TCF1^+^ cells than memory cells (**Figure 6A-B**, **Figure S13**). We also observed that TSCM (CD45RA^+^CD45RO^-^CXCR3^-^CD95^+^CD27^+^) and CXCR3^+^ naïve CD8^+^ T cells express similar levels of TOX, with both populations displaying increased expression of TOX relative to naïve cells (TSCM: p=0.0130, FC=1.21; naïve CXCR3^+^: p=0.0013, FC=1.19) (**Figure 6C**). Interestingly, we also noted increased TCF1 expression on naïve CXCR3^+^ cells from T1D donors relative to control donors (**Figure 6D**) (p=0.0317, FC=1.81). Supporting the notion that these cells possess a transitional phenotype, TCF1^+^ cells expressed significantly increased expression of CD127 as compared to TCF1^-^ cells (p=0.0107, FC=1.32), as well as reduced PD-1 expression (p=0.0003, FC=0.80), and enhanced CD5 expression (p<0.0001, FC=1.85) (**Figure 6E-G**). Moreover, after four hours of stimulation with PMA/ionomycin, we found that naïve CXCR3^+^ cells upregulated IL-2 and TNFα as compared to the remainder of naïve (CXCR3-) cells (IL-2: p=0.0002, FC=3.11; TNFα: p=0.0006, FC=3.12) (**Figure 6H**, **Figure S13**), but had reduced IFNG expression compared to TSCM and memory cells and similar IL-2 and TNFα expression to TSCM cells, indicative of maintained functionality and potentially a transitional phenotype. Importantly, these data support that this subset exhibits features of stemness as well as polyfunctionality.

**Figure 6.**
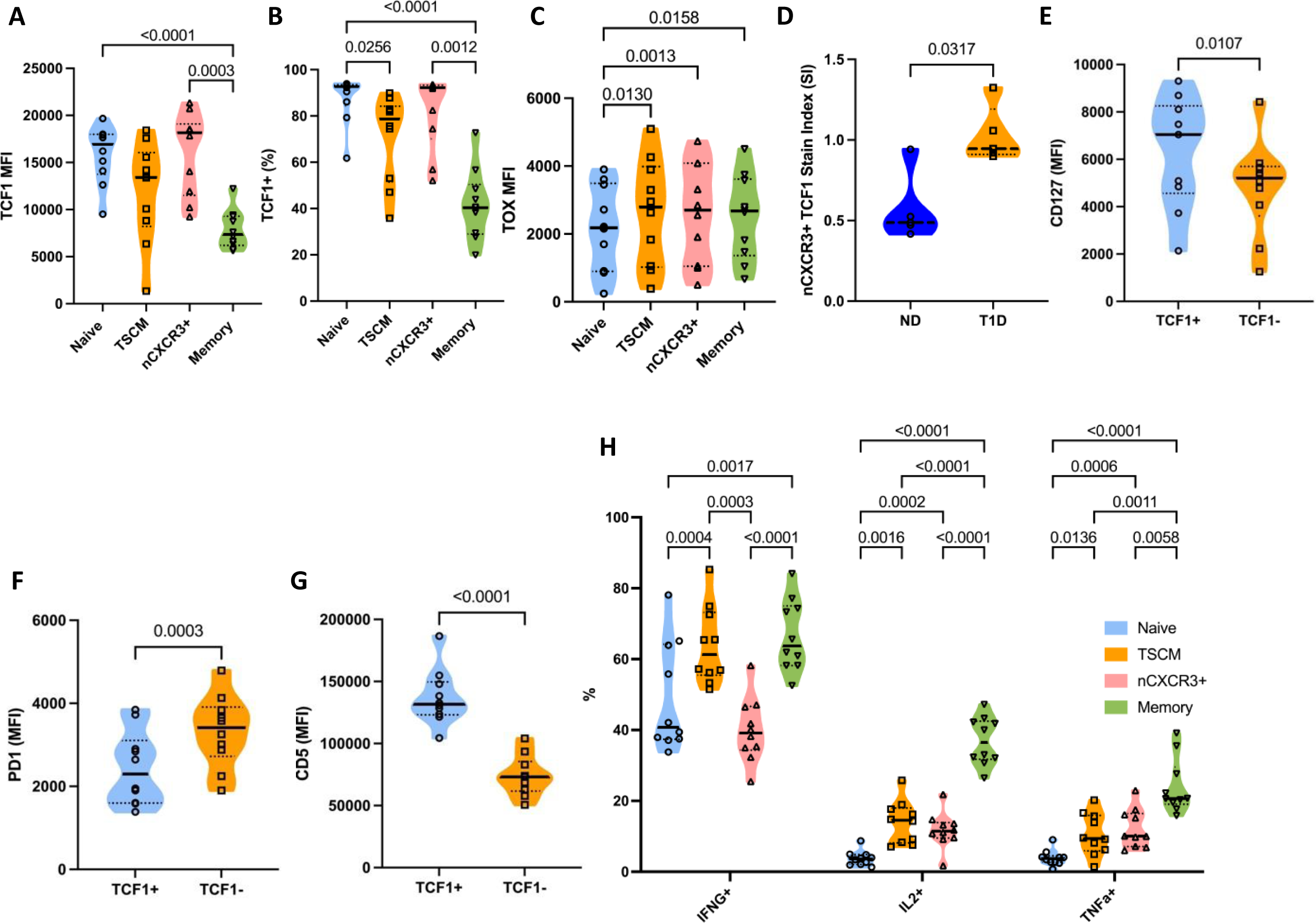
Naïve CXCR3^+^ CD8 T cells possess features of stemness and polyfunctionality. Flow cytometry of cryopreserved pLN cells was performed to assess expression of canonical TSCM markers and cytokine expression. A) Naïve CXCR3^+^ (nCXCR3+) cells resemble TSCM and naïve cells in their expression of TCF1. n=10, p values are the result of a repeated measures one-way ANOVA with Tukey’s multiple comparison test. B) Enrichment of TCF1^+^ cells in naïve CXCR3+ cells as compared to memory cells. N=10, p values are the result of a repeated measures Friedman test with Dunn’s multiple comparison test.C) TSCM and nCXCR3+ cells express more TOX as compared to naïve cells. N=10, p values are the result of a repeated measures one-way ANOVA with Tukey’s multiple comparison test. D) T1D donor nCXCR3+ cells express enhanced TCF1 as quantified by stain index [(median TCF1 expression in nCXCR3+ cells– median TCF1 expression in memory CD8 T cells)/(SD of TCF1 in memory CD8 T cells * 2)] in nCXCR3+ cells from pLN. n=5 per group, p value is the result of a two-tailed mann-whitney test. Upregulation of E) CD127, downregulation of PD-1 (F), and upregulation of CD5 (G) in TCF1^+^ compared to TCF1^-^ cells. N=10, p values are the result of a two-tailed t-test. H) nCXCR3+ cells possess, after 4 hours of PMA/ionomycin stimulation in the presence of gologistop, more IL-2 and TNFα expressing T cells as compared to naive T cells, and more closely resemble TSCM for expression of these molecules. n=10, p values are the result of a repeated measures two-way ANOVA with Tukey’s multiple comparison test.

### TCF1^+^TOX^+^ T cells demonstrate tissue-specific phenotypes and localize to the islets in human T1D

As single cell RNAseq analysis precludes the investigation of cell type localization, we performed PhenoCycler analysis of pancreas and pLN from donor 6551 to examine the tissue localization of naïve-like and terminal effector phenotypes in insulitis (**Figure 7A**). CD3^+^ T cells were classified as TCF1^+^TOX^-^(TCF1^+^, naïve), TCF1^-^TOX^+^ (TOX^+^, memory and exhausted), and TCF1^+^TOX^+^ (activated transitional and effector) populations (**Figure 7B**). TCF1^+^TOX^+^ cells in the pancreas were found more often near or within the islet boundary than TCF1^+^ or TOX^+^ cells, with TOX^+^ cells more proximal than TCF1^+^ (**Figure 7C**). In accordance with our single cell analysis of CD8^+^ T cells across tissues, we observed enhanced inhibitory receptor expression (namely TOX, CD57, and PD-1) in TCF1^+^TOX^+^ and TOX^+^ subsets in the pancreas relative to the pLN (**Figure 7D-E**). Thus, these data indicate a spectrum of memory CD8^+^ T cell phenotypes in the pancreas, which include an activated TCF1^+^TOX^+^ population in close proximity to the islets. Akin to recent reports in NOD mice that precursor diabetogenic T cells are primed and maintained as a self-renewing pool in the pLN and seed terminally differentiated cells that expand in the islets^5^, our data are indicative of distinct tissue programs allowing for maintenance of CD8^+^ T cell populations and prevention of exhaustion in the pLN, as well as acquisition of further effector function in the pancreas during human T1D.

**Figure 7.**
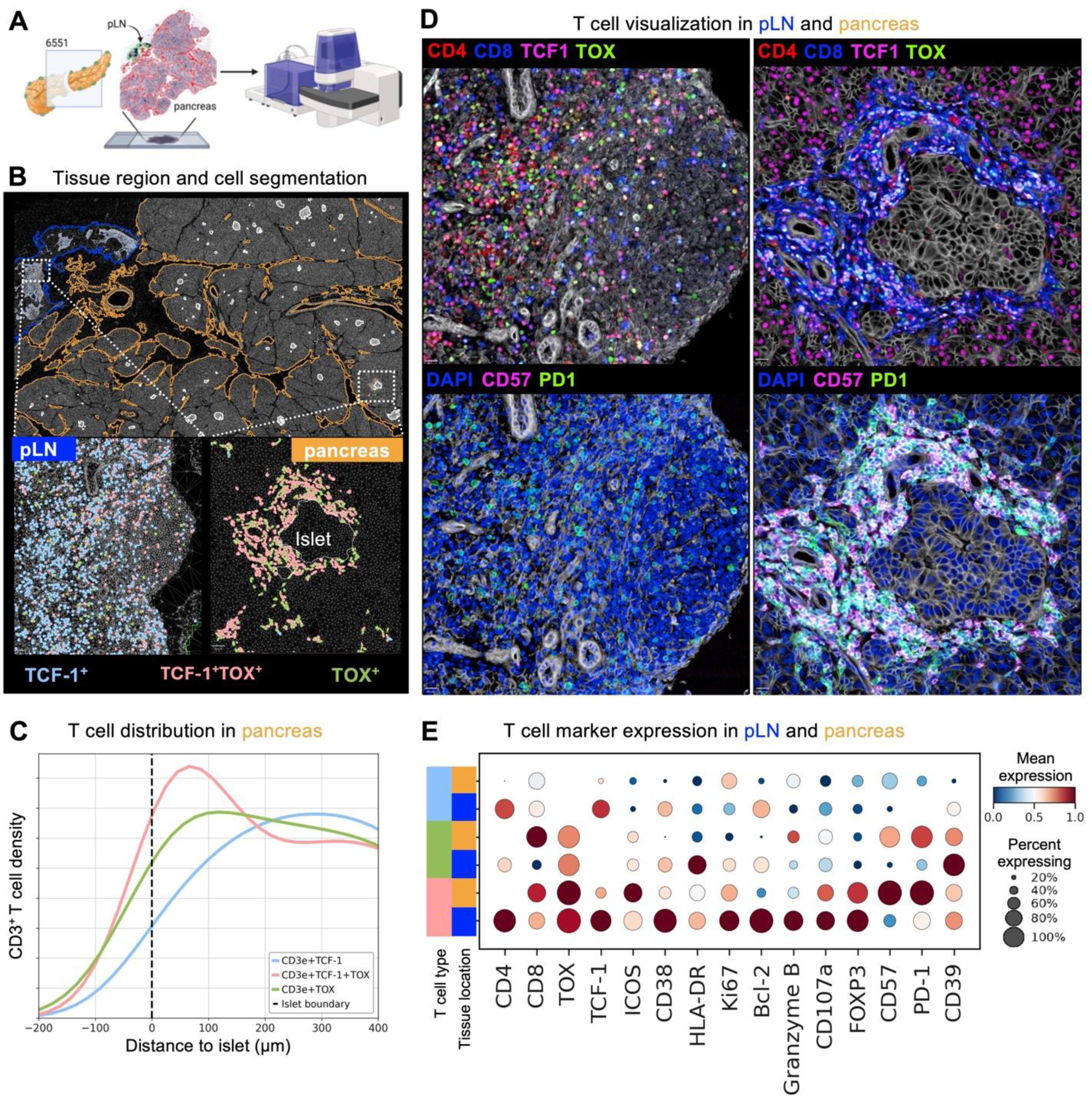
TCF1 and TOX co-expressing populations colocalize to the peri-islet area in human T1D. A) Schematic of phenocycler imaging of embedded pLN and pancreas from nPOD 6551. B) Representative images of pancreas and pLN, with callouts showing TCF1 and TOX expressing populations. C) Line plot illustrating the density of TCF1 and TOX expressing T cell populations across islet distance, with TCF1 and TOX co-expressing cells having greater density nearer to the islets than single positive populations. D) Representative islet images from donor 6551 showing lineage markers (CD4, CD8) as well as TCF1, TOX, CD57 and PD1 alongside nuclear stain DAPI. E) Dotplot showing expression of key memory, negative regulatory and activation related markers across TCF1 and TOX expressing populations.

## Discussion

Despite tremendous progress in characterizing circulating immune cells in T1D, a critical gap in knowledge remains as to alterations in tissue immune phenotypes. Large datasets derived from the pancreas have given insight into disease-related changes in endocrine composition and function^90, 91^, as well as potential changes in immune:islet cell communication^90, 92^, but have limited power to examine immune cell phenotype. Datasets derived from secondary lymphoid organs in T1D are lacking due to the limited availability of samples and, to date, paucity of receptor information^93^. This study begins to address these gaps using complementary methods (i.e., CyTOF, scRNAseq/TCRseq/CITEseq, spatial proteomics) to assess differences in T cell phenotypes and their potential molecular underpinnings within the pLN and pancreas.

CD8^+^ T cells are classically thought to be a major mediator of islet destruction in T1D, given their abundance in the insulitic lesion^94^ and cytolytic function^95–97^. Moreover, recent studies indicated that CD8^+^ T cells with a self-renewing phenotype may be important for T1D pathogenesis, as they are elevated in peripheral blood, and autoreactive cells may be biased towards this phenotype^5, 14, 98^. Our study shows that CD8^+^ T cells expressing markers of a TSCM population (CD45RA, CCR7, CXCR3, CD27, CD28) are increased in T1D pLN. TSCM cells play an important role in memory maintenance and in generating rapidly responding effector cells^99^. It has been shown that serum concentrations of cytokines that promote TSCM development, notably IL-15, are elevated in T1D patients^100, 101^. Moreover, circulating islet antigen-reactive T cells from T1D donors have been shown to possess stem-like features^14^. Enhanced capacity for self-renewal could be clinically detrimental, as multiple therapeutic avenues (e.g., teplizumab, anti-thymocyte globulin [ATG]) have converged on a signature of CD8 T cell exhaustion in therapeutic responders^13, 102^. Our study provides further support for the relevance of this pathway, as IREA analysis revealed an enrichment of IL-15-induced genes across the trajectory of CD8^+^ T cell differentiation in T1D. Interestingly, pathway analysis of naïve cells indicated an enrichment of autophagy and mitophagy in cells from T1D pLN. IL-15 has recently been shown to enable sustained autophagy during activation as compared to inflammatory cytokines^103^, potentially better enabling maintenance of memory populations. We also observed an increase in cytokine signaling and upregulation of the anti-apoptotic factor *BCL2,* which, coupled with the downregulation of multiple exhaustion-associated genes in this cluster in T1D, could suggest that CD8 T cells favor a progenitor exhausted phenotype rather than a terminal exhaustion phenotype in T1D^62^. Further studies are needed to connect IL-15 signaling in T1D to autoreactive T cell stemness and progenitor exhausted phenotypes and to investigate if inhibiting this pathway may aid in establishing durable tolerance. Notably, baricitinib, an inhibitor of Janus Kinase (JAK) 1/2 signaling and thus, common gamma chain cytokine signaling (including IL-15), has recently demonstrated efficacy in preserving β-cell function in recent-onset T1D donors^104^. Interestingly, we observed reduced expression of *IL7R* and *IL2RG* in effector cells derived from the recent-onset T1D pancreas relative to those from paired pLN, indicating that in the pancreas microenvironment, these cells may be more reliant on IL-15 and other effector cytokines, such as IL-21, than IL-2 or IL-7. It remains to be seen whether baricitinib alone could aid in targeting the autoreactive T cell reservoir or if combinatorial therapy will be necessary with T cell exhaustion-inducing therapeutics, such as teplizumab.

Limitations of the study include donor heterogeneity and limited sample size due to the rarity of study tissues. Notably, we were unable to obtain sufficient immune cell numbers from control pancreatic slices to compare immune phenotypes between T1D and ND donors, and our small sampling size from pancreatic slices likely is not representative of the full complement of immune cell phenotypes in the target organ. Our lab is currently utilizing spatial transcriptomic^105^ and proteomic technologies to assess this without the need for immune cell isolation. Moreover, T1D is a heterogeneous disease, and it is possible that immune phenotypes identified through our studies may differ across different age or ancestry groups, which we were not powered to assess. Indeed, though great strides are being made in investigating the genetic drivers of T1D in populations with diverse genetic ancestry^106–108^, there are still many knowledge gaps regarding the importance of known genetic risk loci across groups traditionally underrepresented in biomedical research. Despite this, we believe that the information that we have provided using paired phenotype and receptor information for these subsets can be used by the field to identify CDR3 motifs of further interest.

As it stands, known TCR sequences that correspond to autoreactive T cell specificity are rare^78^. Moreover, matching sequences to those in curated databases such as the Manually curated database of Pathology Associated Sequences (McPAS)^109^ and Immune Epitope Database (IEDB)^110^ introduces a bias toward known specificities and epitopes of interest. Indeed, we were unable to confidently identify a priori T1D antigen autoreactive cells using pMHC dextramer reagents. This could reflect low sampling (10,000 cells per donor), incomplete knowledge of antigen reactivity, and/or individual heterogeneity in autoantigen reactivity. However, utilizing a curated list of experimentally validated T1D clones, we identified 11 putative T1D clones in the pLN based on a priori knowledge. Nevertheless, there is a need to apply new analytical strategies to identify similarities in TCR sequence and transcriptomic features that may indicate a common antigen specificity^111^. We utilized one such tool, CoNGA^31^, to examine joint gene expression and receptor features in T1D. This analysis revealed a T1D-associated enrichment of a TCR cluster comprised of diverse naive CD8^+^ T cells that display features of prior activation (i.e., *KLRK1*, *HCST*, low *CXCR3*), further supporting our gene expression and proteomic data. The feature of low AF4 within this cluster supports previous data wherein autoreactive T cells were found to possess more hydrophobic residues^112^, including within autoreactive receptors detected in the pancreas^79^. Moreover, in T1D pLN we also found enrichment of a more terminally differentiated cluster of CD8^+^ T cells, which displayed expression of cytotoxicity-related genes, *CXCR3*, and memory/exhaustion-related gene *TOX*, while retaining some *TCF7* expression. Though infrequent, we postulated that this population may represent a transitional memory phenotype that is poised to infiltrate the pancreas, as supported by the presence of an expanded putative autoreactive clone in this population in a new-onset T1D donor.

We utilized paired pLN and pancreas T cell data to investigate phenotypes of shared populations and to interrogate whether local expansion of T cells occurred in fresh pancreatic slices. We identified the most abundant shared populations to be effector CD8^+^ T cells, which were the most clonally expanded populations. Moreover, we noted CD8^+^ T cells, as a whole, had reduced expression of genes involved in promoting stem-like/naïve phenotypes (*LEF1, SELL*) in pancreas compared to pLN. This could, in part, be driven by the local inflammatory milieu. Indeed, we observed enrichment of multiple inflammatory cytokine signaling pathways, namely IL-6 and TNFα, in pancreas relative to pLN T cells, while we observed upregulation of Myc, mTORC1, and Wnt signaling pathways in pLN relative to pancreas T cells. While sharing of TCR clones was limited between donors and difficult to detect due to the sparsity of single cell data, we were able to detect intra-donor sharing of clones across tissues and between clusters. Donor 6551, a 20-year-old donor with 0.58 years T1D duration, particularly notable for insulitis severity, possessed expanded CD8^+^ Teff clones that were shared between tissues and between clusters. In the longer duration (4 years) donor with less insulitis (6536), we were able to detect sharing of only one effector CD8^+^ clone across clusters and one CD4^+^ clone across tissues.

Using multiplex spatial proteomic profiling, we were able to show that in the same T1D donor with marked insulitis, case 6551, TCF1 and TOX expressing T cell populations can be found in pLN and pancreas. Importantly, we found that TCF1^+^TOX^+^ cells possess a more terminal effector phenotype in the pancreas and can be found proximal to the islets. We also found by flow cytometry that pLN naive CXCR3^+^ T cells have similar expression of transcription factors TCF1 and TOX to TSCM cells, and produce similar levels of proinflammatory cytokines upon activation. Akin to human peripheral blood data, we noted that TCF1-expressing populations in the pLN tended to express less PD-1 and more CD127^113^ and CD5^114^, indicative of early differentiation stage and activation potential. In accordance with murine studies^5^, these data collectively support that T cell clones are supplied from the pLN and adopt a more effector and terminally differentiated phenotype in the pancreas during human T1D development. Previously, our lab reported increased circulating CXCR3^lo^CD8^+^ T cells in T1D^75^ following phenotyping of a cohort of over 824 cross-sectional peripheral blood samples. The present study supports a role for these cells in the draining lymph node: a small set of related terminal effector CXCR3-expressing cells could be found in the pLN and a larger fraction in the target organ, suggesting that these cells share some features of stemness (TCF1 expression) and are more polyfunctional as compared to naïve cells. Identifying subsets of interest that are relevant in both peripheral blood and the target organ could aid in efforts to selectively inhibit autoreactivity and monitor outcomes. Importantly, combination CXCR3 antagonism with ACT-777991 and α-CD3 treatment leads to disease remission in the NOD mouse^115^, and first-in-human trials in healthy adults have shown this compound to be tolerable^116^. Thus, combinatorial strategies to induce tolerogenic phenotypes while targeting pathways relevant to tissue immune phenotypes may represent an avenue to achieve durable tolerance. Taken together, our studies indicate that memory and effector T cell programs in T1D are maintained in the pLN and enhanced locally in the pancreas, likely driven by aberrant regulation of cytokine signaling and impacted by genotype at T1D risk loci. Future directions validating the functionality of clones identified herein and investigating the cellular interactions of CXCR3^+^CD8^+^ T cells across the differentiation trajectory will inform the development of therapeutics to modulate them, providing novel avenues to delay or prevent T1D progression.

## Supporting information

Supplemental Values

## Author Contributions

LDP: Designing research studies, conducting experiments, acquiring data, analyzing data, writing the manuscript (original draft), funding acquisition

HRS: Conducting experiments, acquiring data, writing the manuscript (reviewing and editing)

JS: Conducting experiments, acquiring data, analyzing data, writing the manuscript (original draft) ALP: Writing the manuscript (reviewing and editing)

RLB: Acquiring data, writing the manuscript (reviewing and editing) CHW: Writing the manuscript (reviewing and editing)

MAA: Writing the manuscript (reviewing and editing), funding acquisition

RB: Analyzing data, writing the manuscript (reviewing and editing), interpreting data

MAB: Designing research studies, interpreting data, writing the manuscript (reviewing and editing), project oversight

TMB: Designing research studies, interpreting data, writing the manuscript (reviewing and editing), funding acquisition, project oversight

## Acknowledgments

We thank the nPOD donor families for their important contributions to this research. We also thank the nPOD Organ Processing and Pathology Core (OPPC) team, namely Irina Kusmartseva, Helmut Hiller, and Maria Beery, for their efforts in case acquisition, processing, and distribution.

