## Supplementary Figures and Tables for "Aberrant immune regulation and enrichment of stem-like CD8^+^ T cells in the pancreatic lymph node during type 1 diabetes development"

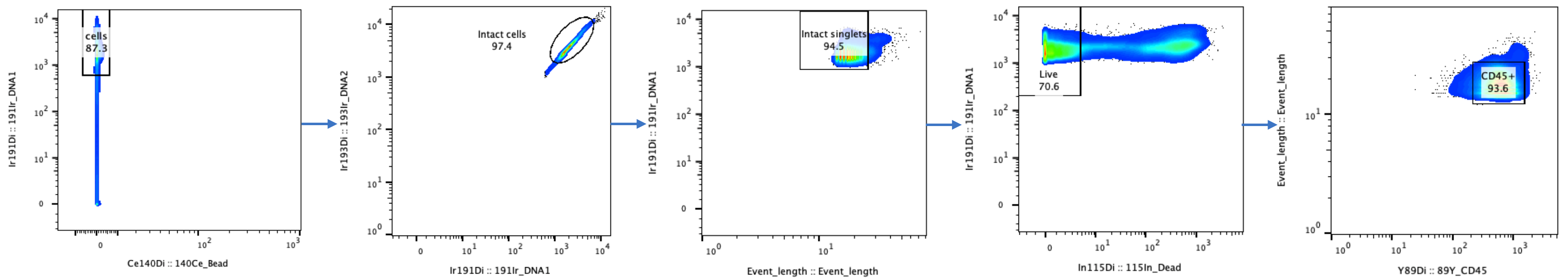

cells
87.3
Ir191Di :: 191Ir_DNA1
Ce140Di :: 140Ce_Bead
Intact cells
97.4
Ir193Di :: 193Ir_DNA2
Ir191Di :: 191Ir_DNA1
Intact singlets
94.5
Ir191Di :: 191Ir_DNA1
Event_length :: Event_length
Live
70.6
Ir191Di :: 191Ir_DNA1
In115Di :: 115In_Dead
CD45+
93.6
Event_length :: Event_length
Y89Di :: 89Y_CD45

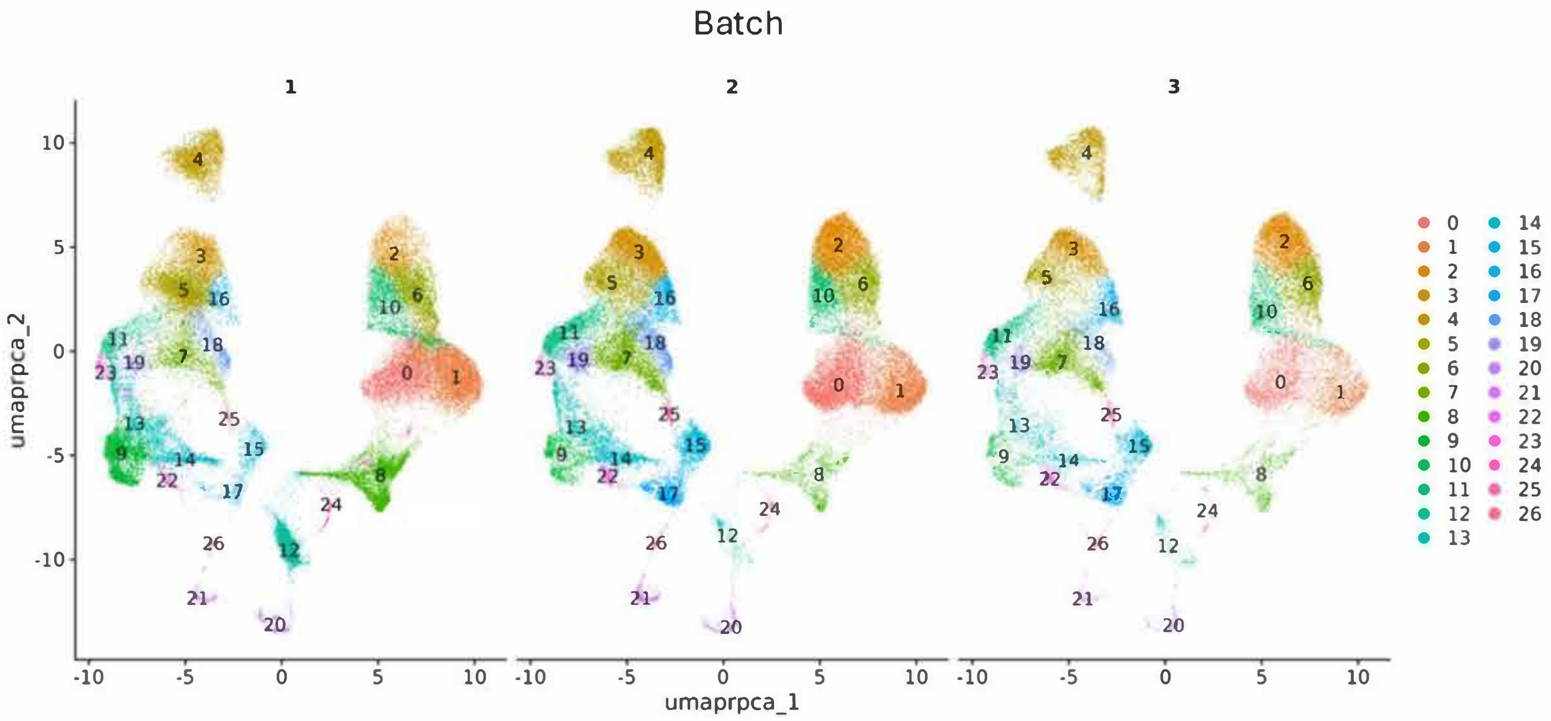
Batch
1
2
3
umaprpca_2
umaprpca_1
0 1 2 3 4 5 6 7 8 9 10 11 12 13 14 15 16 17 18 19 20 21 22 23 24 25 26

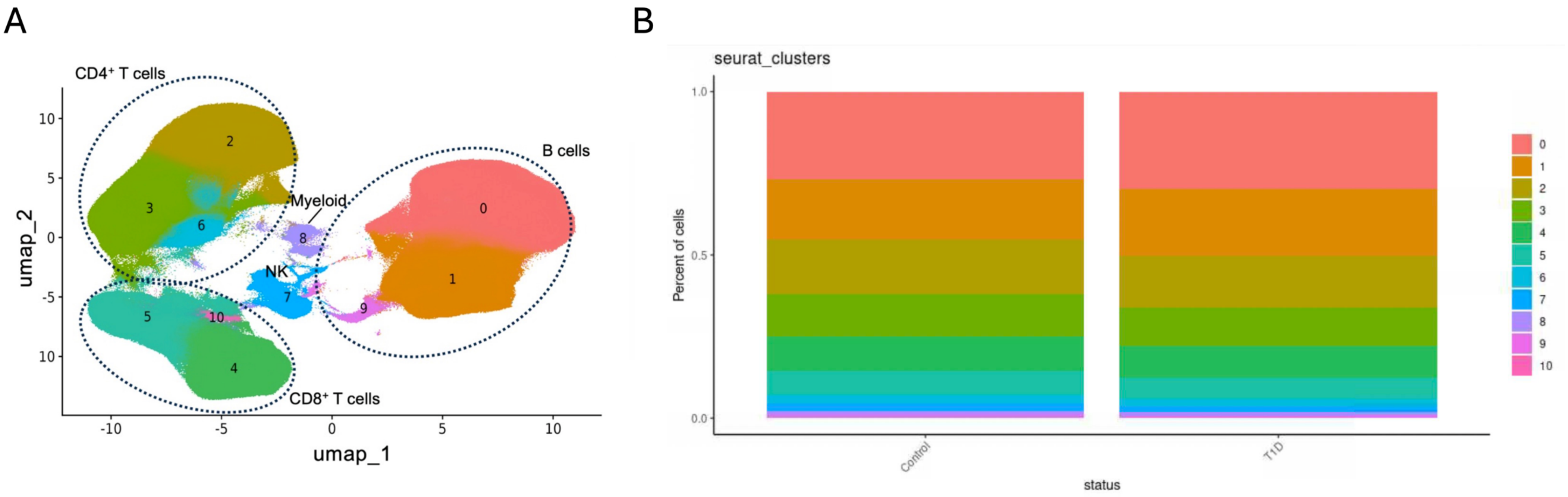
A
CD4⁺ T cells
B cells
Myeloid
NK
CD8⁺ T cells
umap_1
umap_2
B
seurat_clusters
Percent of cells
status
Control
T1D

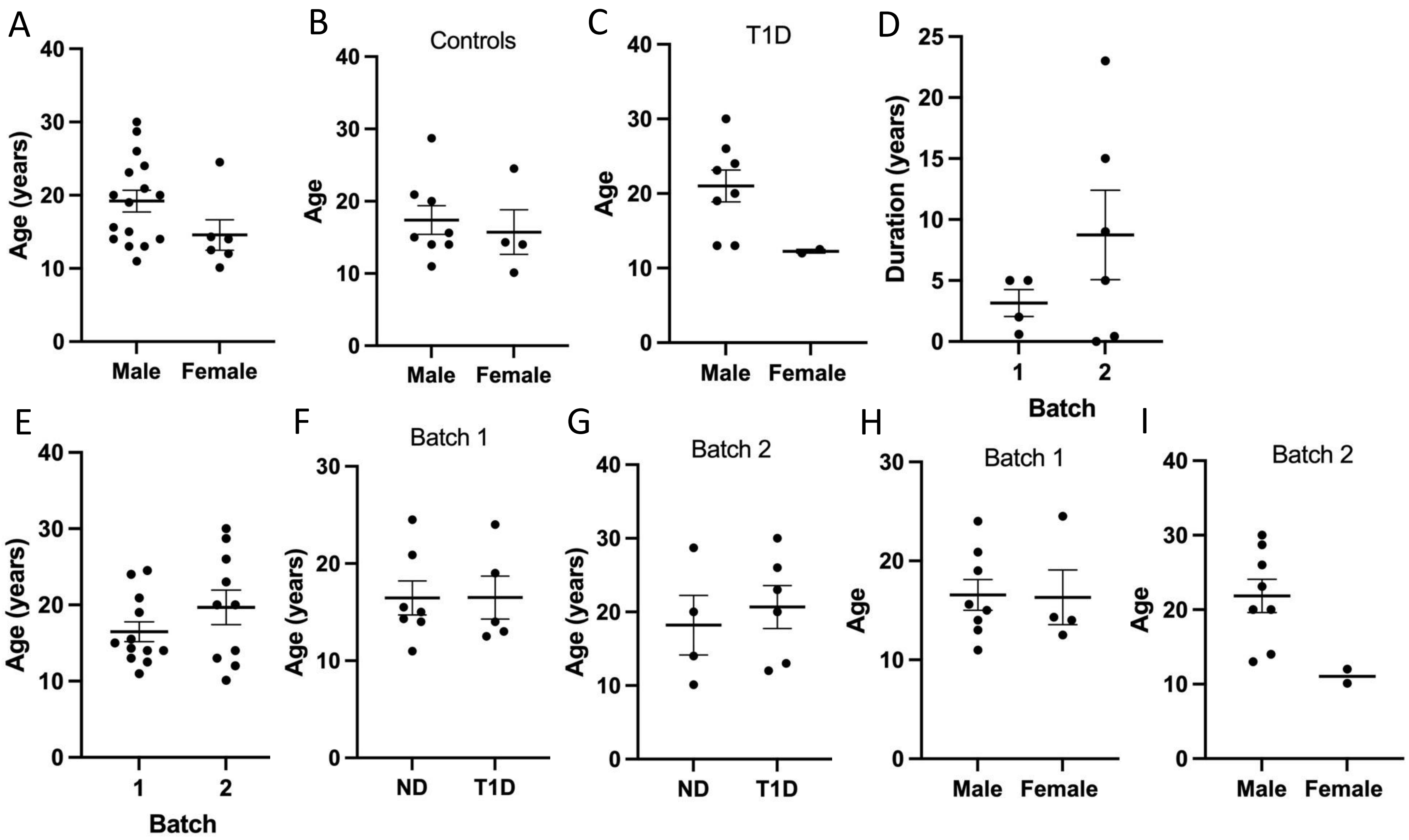
A
Age (years)
Male
Female
B
Controls
Age
Male
Female
C
T1D
Age
Male
Female
D
Duration (years)
1
2
Batch
E
Age (years)
1
2
Batch
F
Batch 1
Age (years)
ND
T1D
G
Batch 2
Age (years)
ND
T1D
H
Batch 1
Age
Male
Female
I
Batch 2
Age
Male
Female

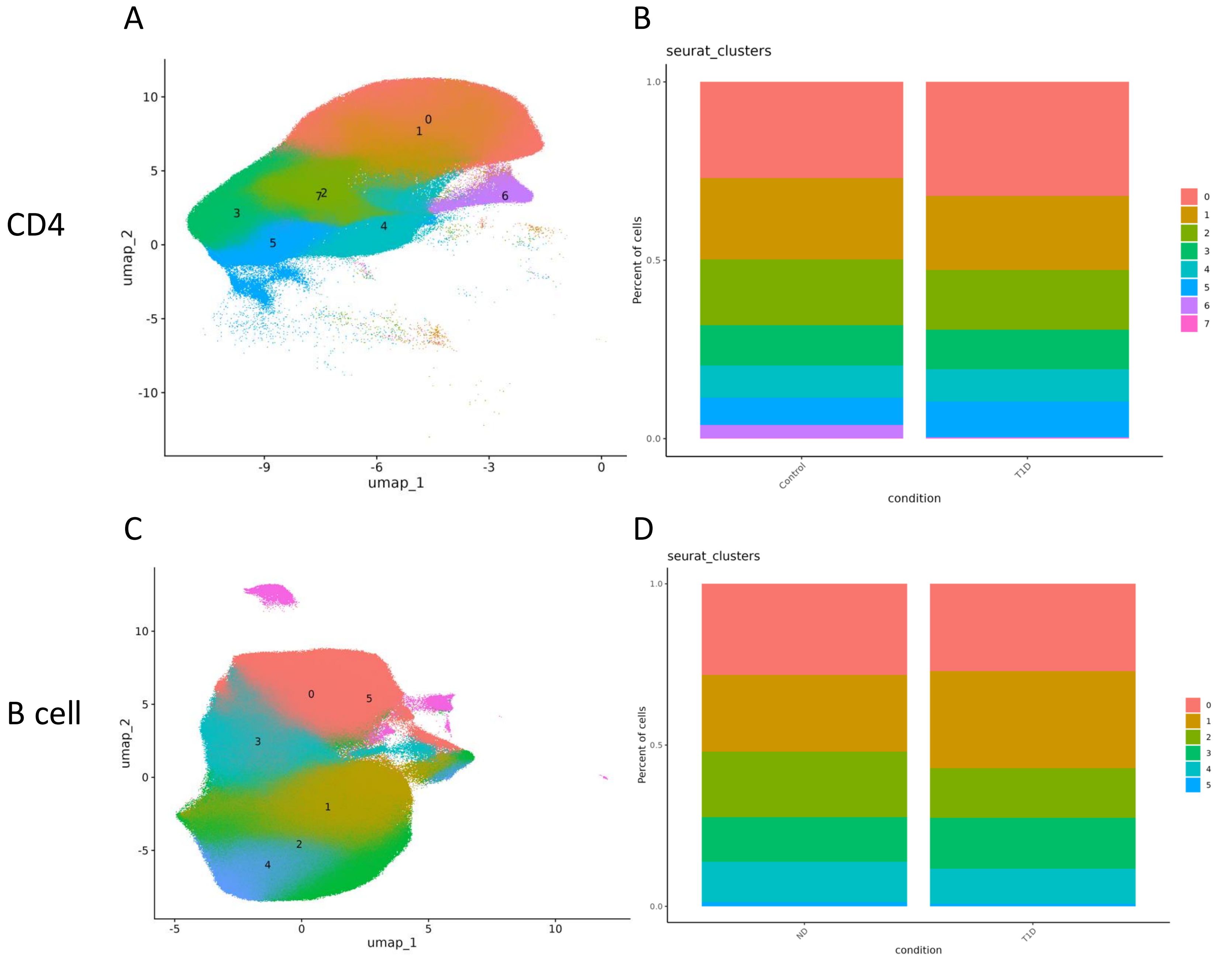
A
B
CD4
seurat_clusters
umap_1
umap_2
Percent of cells
condition
Control
T1D
C
D
B cell
seurat_clusters
umap_1
umap_2
Percent of cells
condition
ND
T1D

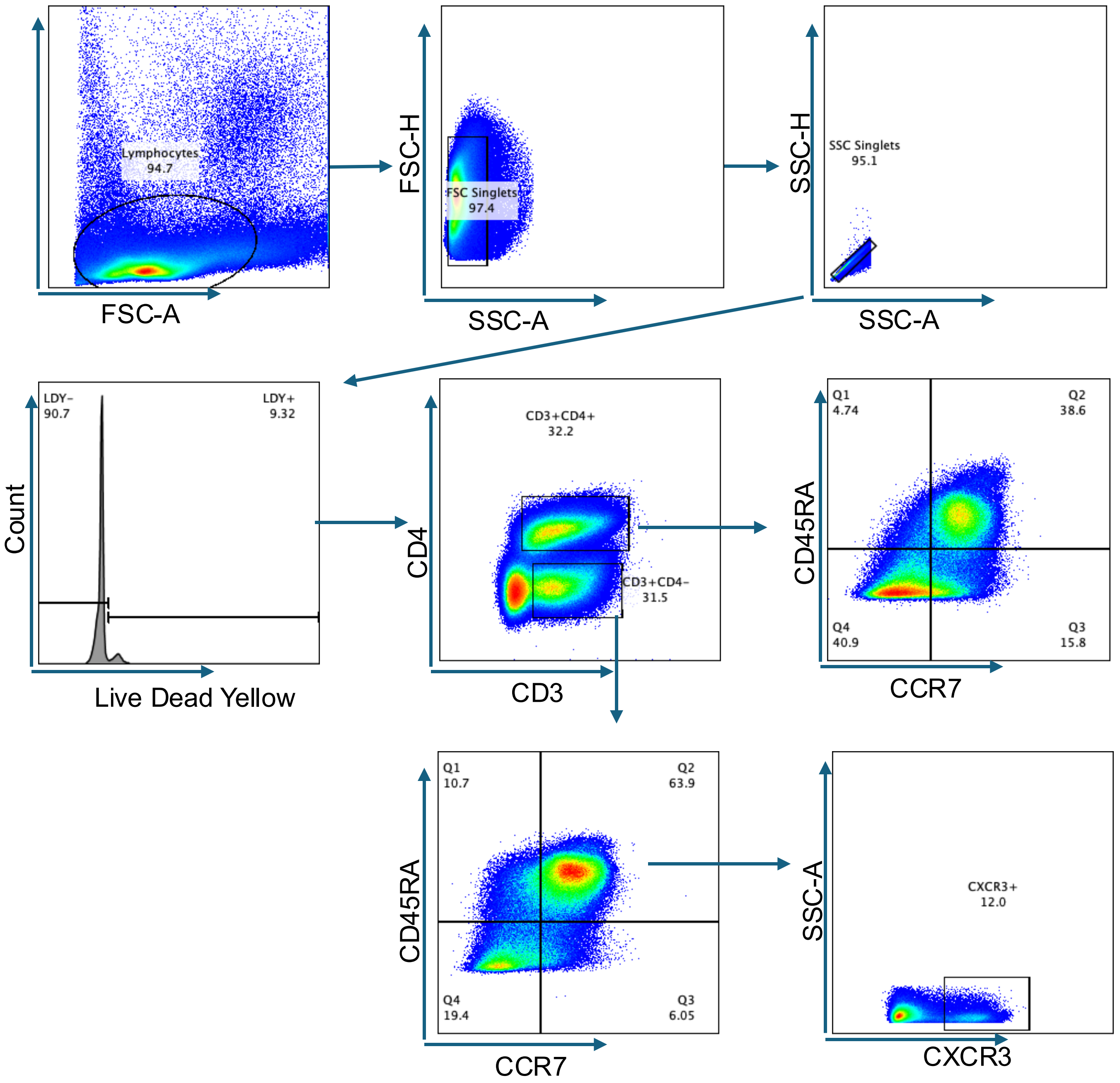
Lymphocytes
94.7
FSC-A
FSC-H
FSC Singlets
97.4
SSC-A
SSC-H
SSC Singlets
95.1
SSC-A
LDY-
90.7
LDY+
9.32
Count
Live Dead Yellow
CD3+CD4+
32.2
CD3+CD4-
31.5
CD4
CD3
Q1
4.74
Q2
38.6
Q4
40.9
Q3
15.8
CD45RA
CCR7
Q1
10.7
Q2
63.9
Q4
19.4
Q3
6.05
CD45RA
CCR7
CXCR3+
12.0
SSC-A
CXCR3

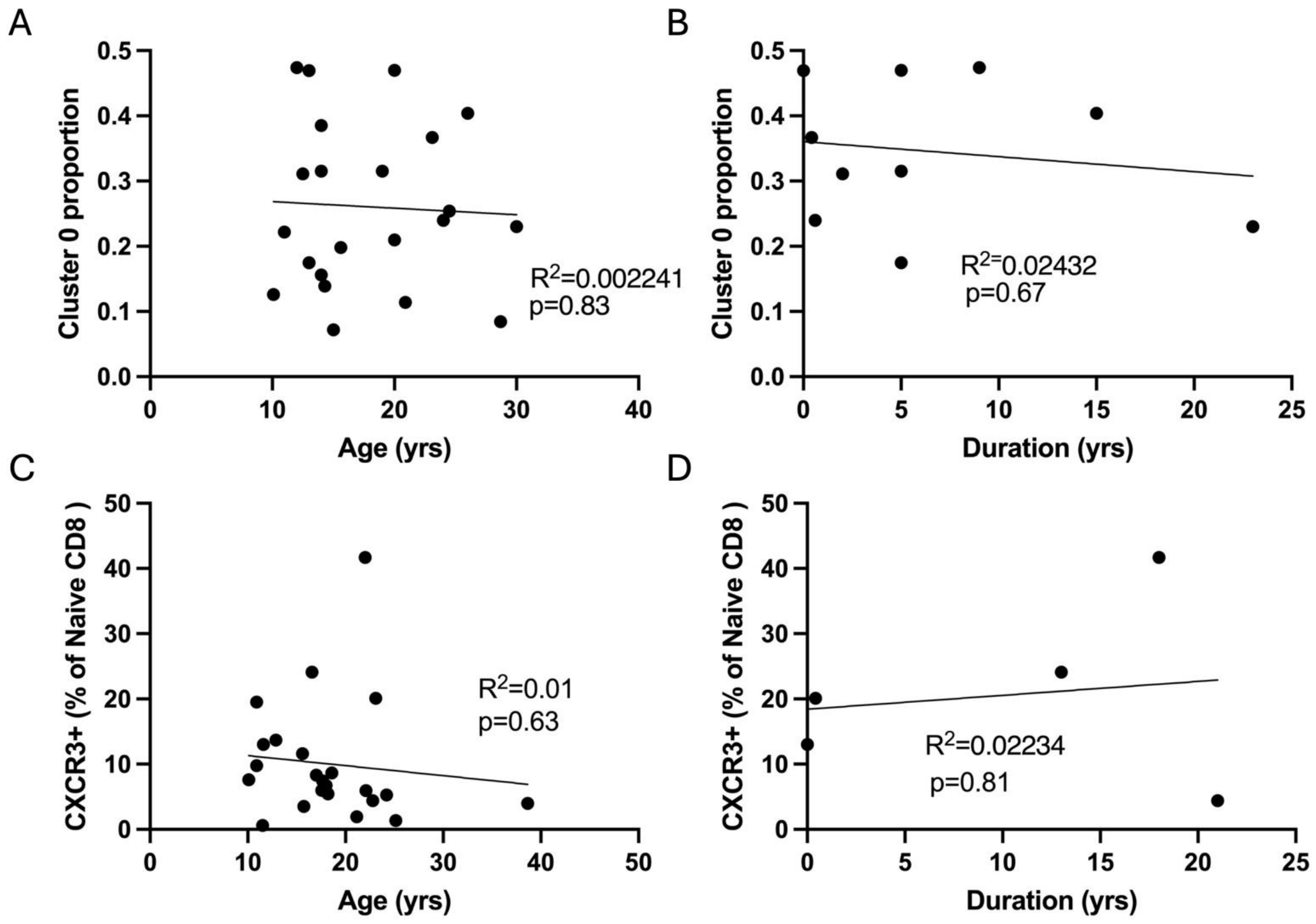
A
Cluster 0 proportion
Age (yrs)
R2=0.002241
p=0.83
B
Cluster 0 proportion
Duration (yrs)
R2=0.02432
p=0.67
C
CXCR3+ (% of Naive CD8 )
Age (yrs)
R2=0.01
p=0.63
D
CXCR3+ (% of Naive CD8 )
Duration (yrs)
R2=0.02234
p=0.81

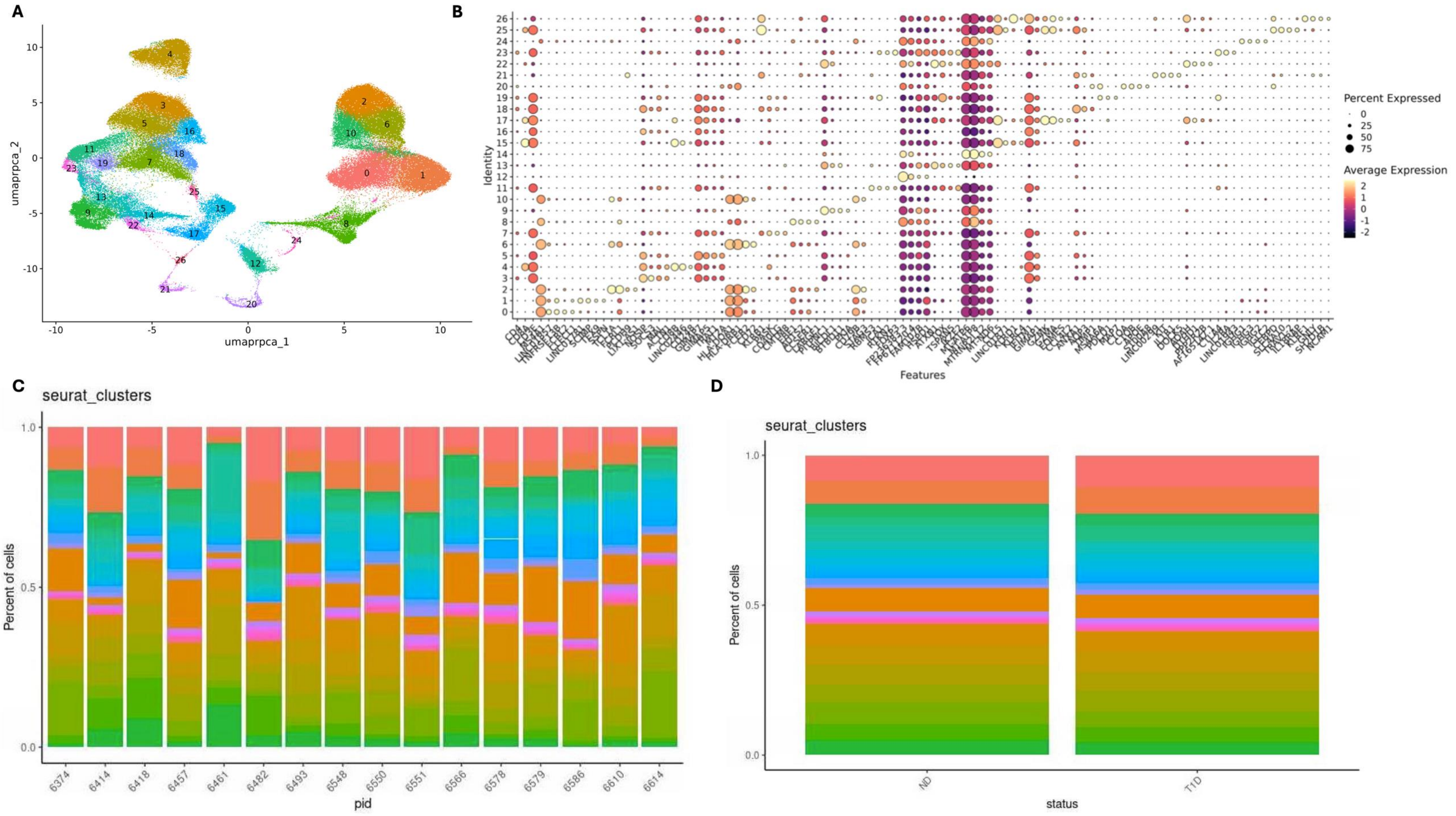
A
B
C
D
umaprpca_1
umaprpca_2
Identity
Features
Percent Expressed
0
25
50
75
Average Expression
seurat_clusters
Percent of cells
pid
6374
6414
6418
6457
6461
6482
6493
6548
6550
6551
6566
6578
6579
6586
6610
6614
seurat_clusters
Percent of cells
status
ND
T1D

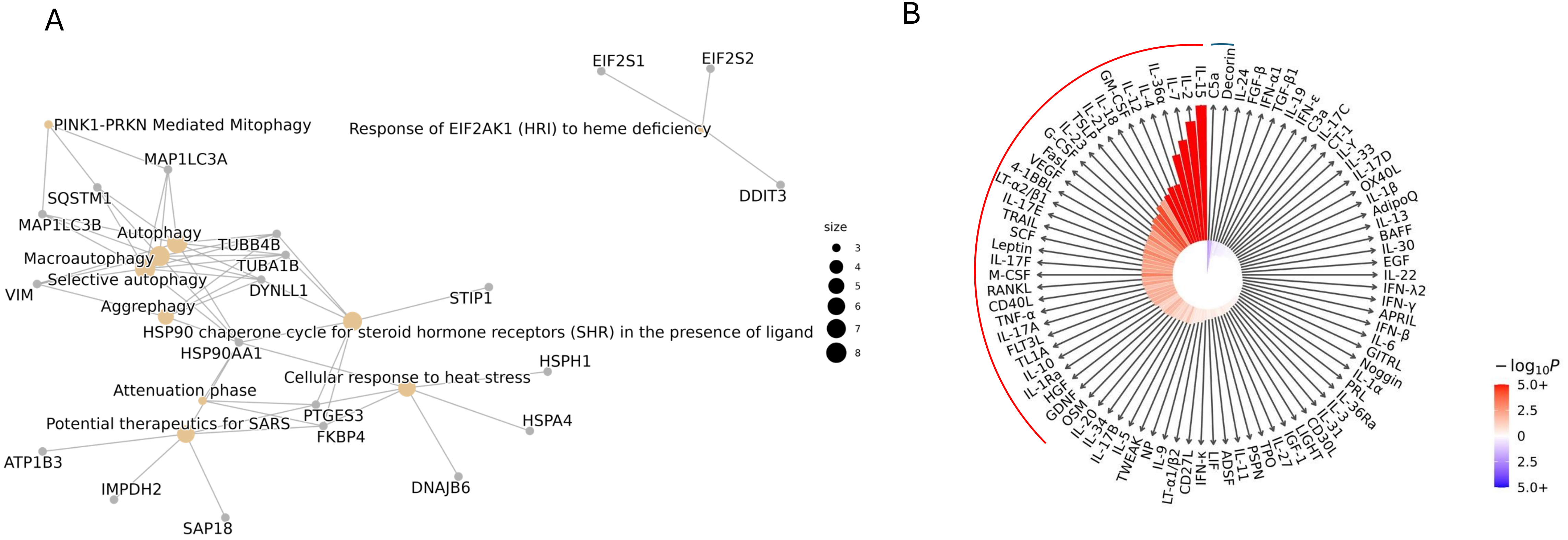
A
PINK1-PRKN Mediated Mitophagy
MAP1LC3A
SQSTM1
MAP1LC3B
Autophagy
TUBB4B
Macroautophagy
TUBA1B
Selective autophagy
VIM
DYNLL1
Aggrephagy
STIP1
HSP90 chaperone cycle for steroid hormone receptors (SHR) in the presence of ligand
HSP90AA1
HSPH1
Cellular response to heat stress
Attenuation phase
PTGES3
Potential therapeutics for SARS
HSPA4
FKBP4
ATP1B3
IMPDH2
DNAJB6
SAP18
EIF2S1
EIF2S2
Response of EIF2AK1 (HRI) to heme deficiency
DDIT3
size
3
4
5
6
7
8
B
IL-15
C5a
Decorin
IL-24
FGF-β
IFN-α1
TGF-β1
IL-19
IFN-ε
C3a
IL-17C
CT-1
IL-33
IL-17D
OX40L
IL-1β
AdipoQ
IL-13
BAFF
IL-30
EGF
IL-22
IFN-λ2
IFN-γ
APRIL
IFN-β
IL-6
GITRL
Noggin
IL-1α
PRL
IL-36Ra
IL-31
CD30L
LIGHT
IGF-1
IL-27
TPO
PSPN
IL-11
ADSF
LIF
IFN-κ
CD27L
LT-α1/β2
IL-9
NP
TWEAK
IL-5
IL-17B
IL-34
IL-20
OSM
GDNF
HGF
IL-1Ra
IL-10
TL1A
FLT3L
IL-17A
TNF-α
CD40L
RANKL
M-CSF
IL-17F
Leptin
SCF
TRAIL
IL-17E
LT-α2/β1
4-1BBL
VEGF
FasL
G-CSF
IL-23
TSLP
IL-21
IL-18
GM-CSF
IL-12
IL-4
IL-36α
IL-7
IL-2
$-\log_{10}P$
5.0+
2.5
0
2.5
5.0+

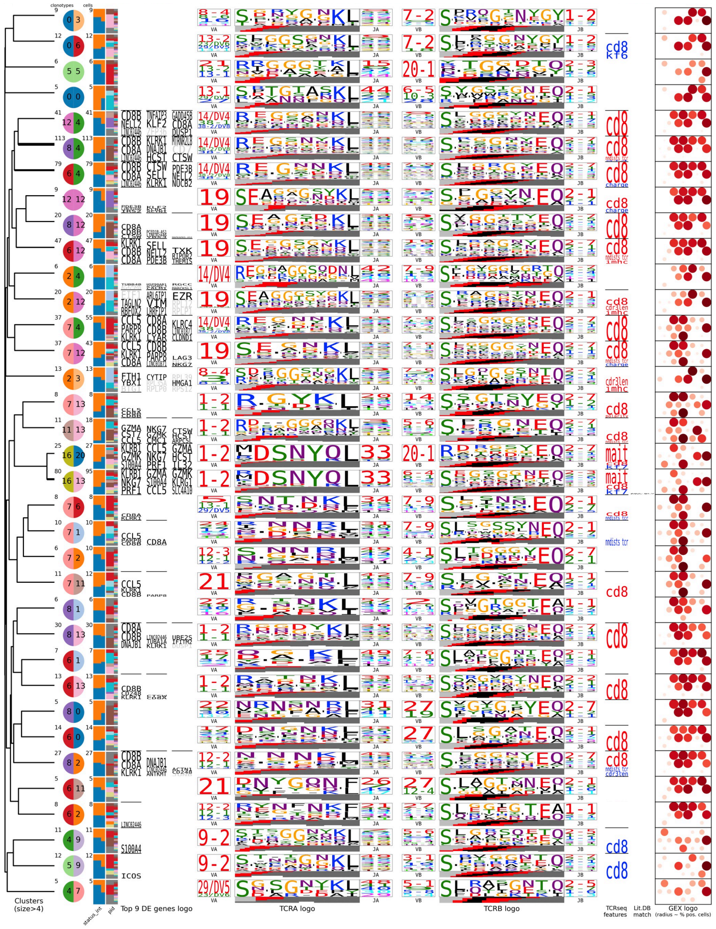

clonotypes
cells
Clusters (size>4)
Top 9 DE genes logo
TCRA logo
TCRB logo
TCRseq features
Lit.DB match
GEX logo (radius ~ % pos. cells)

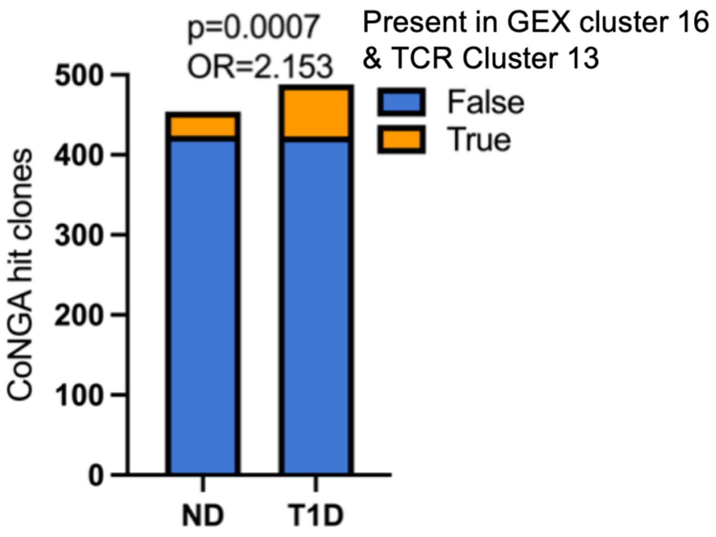
p=0.0007
OR=2.153
Present in GEX cluster 16
& TCR Cluster 13
False
True
CoNGA hit clones
500
400
300
200
100
0
ND
T1D

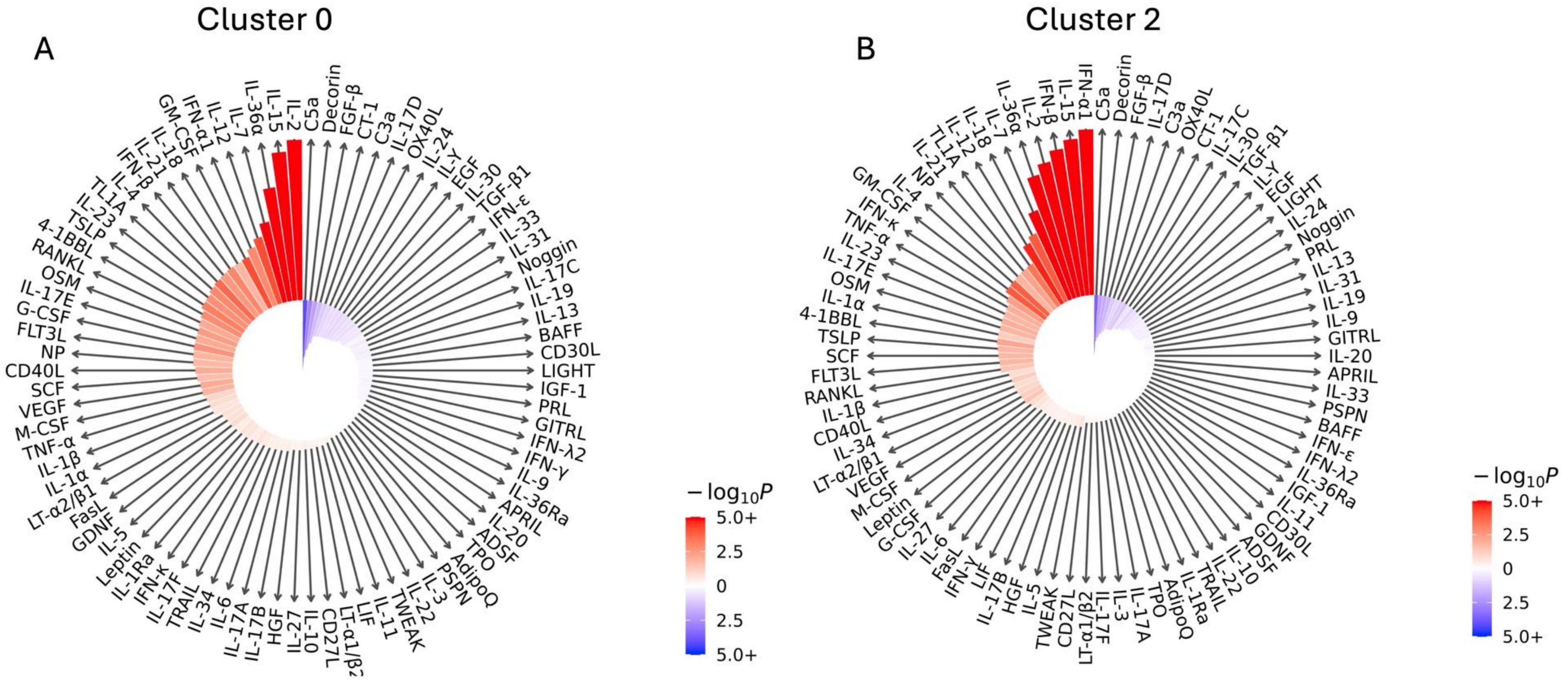

Cluster 0
A
IL-2
IL-15
IL-36α
IL-7
IL-12
IFN-α1
GM-CSF
IL-18
C5a
Decorin
FGF-β
CT-1
C3a
IL-17D
OX40L
IL-24
IL-Y
EGF
IL-30
TGF-β1
IFN-ε
IL-33
IL-31
Noggin
IL-17C
IL-19
IL-13
BAFF
CD30L
LIGHT
IGF-1
PRL
GITRL
IFN-λ2
IFN-γ
IL-9
IL-36Ra
APRIL
IL-20
ADSF
TPO
AdipoQ
PSPN
IL-3
IL-22
TWEAK
IL-11
LIF
LT-α1/β2
CD27L
IL-10
IL-27
HGF
IL-17B
IL-17A
IL-6
IL-34
TRAIL
IL-17F
IFN-κ
IL-1Ra
Leptin
IL-5
GDNF
FasL
LT-α2/β1
IL-1α
IL-1β
TNF-α
M-CSF
VEGF
SCF
CD40L
NP
FLT3L
G-CSF
IL-17E
OSM
RANKL
4-1BBL
TSLP
$-\log_{10}P$
5.0+
2.5
0
2.5
5.0+
Cluster 2
B
IFN-α1
IL-15
IFN-β
IL-2
IL-36α
IL-7
IL-18
IL-12
C5a
Decorin
FGF-β
IL-17D
C3a
OX40L
CT-1
IL-17C
IL-30
TGF-β1
IL-Y
EGF
LIGHT
IL-24
Noggin
PRL
IL-13
IL-31
IL-19
IL-9
GITRL
IL-20
APRIL
IL-33
PSPN
BAFF
IFN-ε
IFN-λ2
IL-36Ra
IGF-1
IL-11
CD30L
GDNF
ADSF
IL-10
IL-22
TRAIL
IL-1Ra
AdipoQ
TPO
IL-17A
IL-3
IL-17F
LT-α1/β2
CD27L
TWEAK
IL-5
HGF
IL-17B
LIF
IFN-γ
FasL
IL-6
IL-27
G-CSF
Leptin
M-CSF
VEGF
LT-α2/β1
IL-34
CD40L
IL-1β
RANKL
FLT3L
SCF
TSLP
4-1BBL
IL-1α
OSM
IL-17E
IL-23
TNF-α
IFN-κ
GM-CSF
$-\log_{10}P$
5.0+
2.5
0
2.5
5.0+

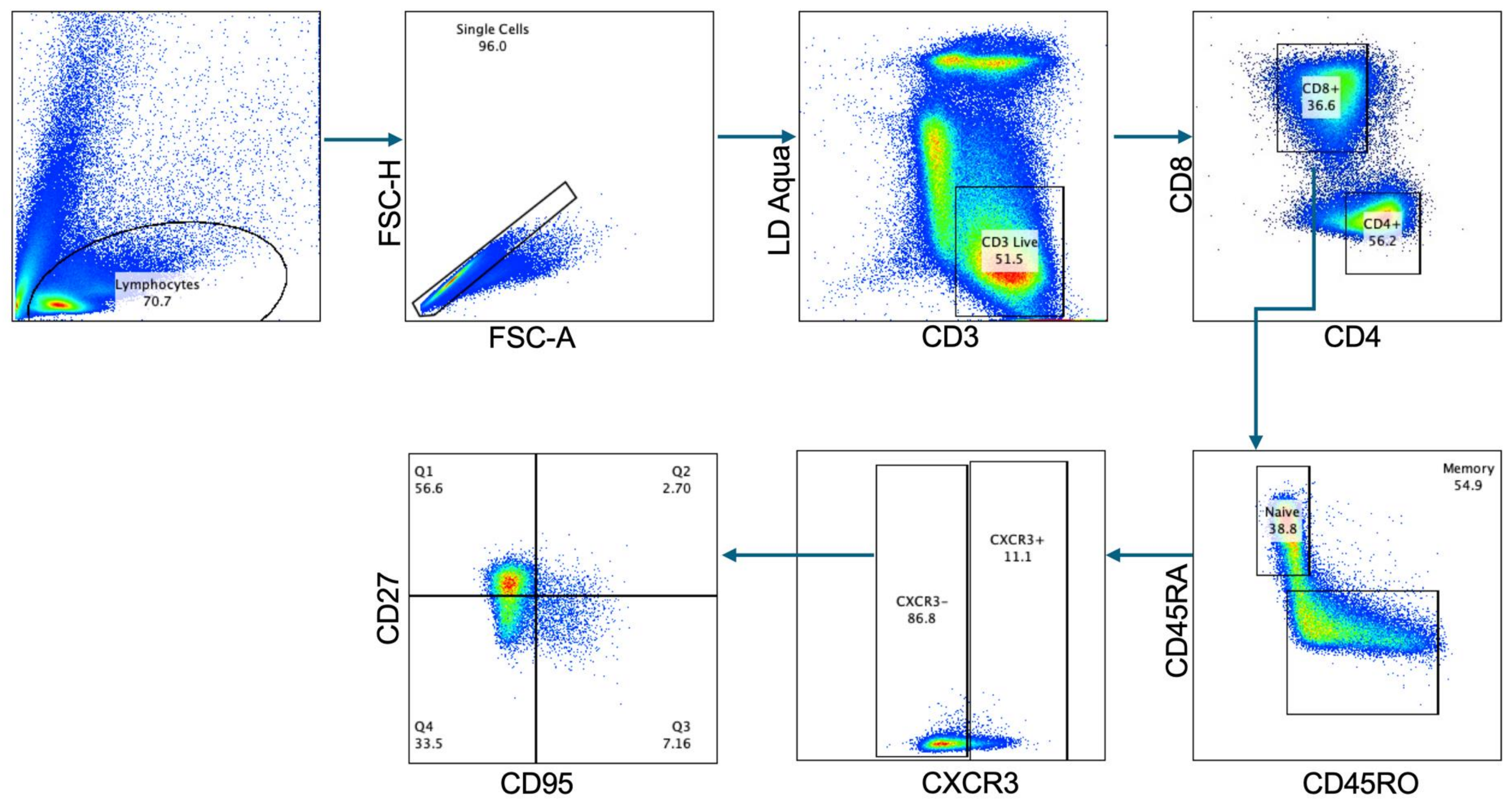

Lymphocytes
70.7
Single Cells
96.0
FSC-H
FSC-A
CD3 Live
51.5
LD Aqua
CD3
CD8+
36.6
CD4+
56.2
CD8
CD4
Memory
54.9
Naive
38.8
CD45RA
CD45RO
CXCR3+
11.1
CXCR3-
86.8
CXCR3
Q1
56.6
Q2
2.70
Q4
33.5
Q3
7.16
CD27
CD95

**Figure S1. Gating scheme for CyTOF data**. Cells were gated from any residual beads after bead removal with premessa, followed by intact cells, intact singlets, live cells, and $CD45^+$ cells.

**Figure S2. No significant differences in pLN cluster frequency across technical batches.** UMAP of pLN scRNAseq data split by batch.

**Figure S3. Similar profiles of total immune cells in the pLN between ND and T1D donors by CyTOF.** A) UMAP illustrating clustering of total $CD45^+$ pLN cells. B) Barplot of CyTOF clusters split by clinical status (n=12 ND, n=10 T1D) shows similar composition of immune cell subsets.

**Figure S4. Covariate information in CyTOF batches.** A) Age of the entire cohort separated by biological sex. B) Age of only controls in the cohort separated by biological sex. C) Age of only T1D donors in the cohort separated by biological sex. D) T1D duration separated by batch. E) Age of the entire cohort separated by batch. F) Age of donors in batch 1 separated by clinical status. G) Age of donors in batch 2 separated by clinical status. H) Age of donors in batch 1 separated by biological sex. I) Age of donors in batch 2 separated by biological sex. All nonsignificant results after performing either two-tailed T test or Mann Whitney test.

**Figure S5. Similar frequency of CD4 and B cell CyTOF clusters by clinical status.** A) UMAP of CD4 T cell CyTOF data. B) Bar plot of clusters from A separated by clinical status (n=12 ND, n=10 T1D). C) UMAP of B cell CyTOF data. D) Bar plot of clusters from C separated by clinical status.

**Figure S6. Gating scheme for assessing $CXCR3^+$ T cells in pLN by flow cytometry.** Lymphocytes were gated to exclude doublets and dead cells, followed by discrimination of $CD3^+CD4^+$ and $CD3^+CD4^-$ (i.e., $CD8^+$) T cells. $CD3^+CD4^+$ and $CD3^+CD4^-$ subsets were gated for naïve markers CD45RA and CCR7, and within the $CD3^+CD4^-CD45RA^+CCR7^+$ population, $CXCR3^+$ cells were gated.

**Figure S7. No significant association of naïve $CXCR3^+$ subsets with disease duration or age.** A) Correlation of CyTOF cluster 0 ($CD8^+CD45RA^+CCR7^+CD27^+CD28^+CXCR3^+$) proportion with age in all donors. B) Correlation of CyTOF cluster 0 proportion with disease duration in T1D donors only. C) Correlation of flow cytometry $CXCR3^+$ naïve CD8 T cell population with age in all donors. D) Correlation of flow cytometry $CXCR3^+$ naïve CD8 T cell population with disease duration in only T1D donors. P and $R^2$ values are the result of Pearson correlation.

**Figure S8. No significant changes in scRNAseq clusters by clinical status.** A) UMAP of pLN scRNAseq data. B) Dot plot showing average expression and percent expression of marker genes per cluster. C) Bar plot of cluster proportion per donor. D) Bar plot of cluster proportion by clinical status.

**S9. Naïve T cell pathway and immune response enrichment analysis (IREA) analysis.** A) Reactome analysis of naïve CD8 T cells indicates upregulation of autophagy and mitophagy in T1D pLN. B) IREA analysis of naïve CD8 T cells implicates IL-15 as a driver of altered phenotypes.

Differentially expressed genes of $p<0.05$ and $Log2FC>0.26$ were used as input into these analyses. Pathway analysis was performed with the ReactomePA R package, and results are reported when $p<0.05$. Bar height in the compass plot of IREA results is indicative of enrichment score, while color corresponds to FDR-adjusted P value (two-sided Wilcoxon rank-sum test). Red bar indicates which pathways were significantly upregulated, while the blue bar indicates the pathways that were significantly downregulated in this cluster.

**Figure S10. CoNGA hits with clone size >4 in pLN.** Semicircles indicate gene expression and TCR cluster, from left to right. Bar graphs illustrate the proportion of the cluster per status (leftmost bar) and dataset per status (rightmost bar). Orange=T1D, Blue=ND. Differentially expressed genes from each cluster are shown in the logo plot. TCRa and TCRb gene information, CDR3a and CDR3b logos are provided per cluster. TCR features are also displayed.

**Figure S11 Enriched cluster of MAIT T cells in T1D pLN.** Bar plot showing number of conga hit clones falling into gene expression (GEX) cluster 16 and TCR cluster 13, which possesses MAIT T cell gene expression and TCR features, by clinical status. P-value is the result of a two-way Fisher's exact test.

**Figure S12. IREA Analysis of pLN vs pancreas CD8 T cells**. A) Compass plot of IREA results using the top genes upregulated in pancreas $CD8^+$ cluster 0 cells ($p<0.05$, $log2FC>0.26$) as input. B) Compass plot of IREA results using the top genes upregulated in pancreas $CD8^+$ cluster 2 cells ($p<0.05$, $log2FC>0.26$) as input. Bar height is indicative of enrichment score, while color corresponds to FDR-adjusted P value (two-sided Wilcoxon rank-sum test).

**Figure S13. Gating scheme for panel assessing stemness features.** Flow cytometry of cryopreserved pLN cells was performed to assess expression of canonical Tscm markers and cytokine expression after activation for 4 hours in the presence of PMA/ionomycin and GolgiStop. Single lymphocytes were gated prior to live $CD3^+$ cells, and CD4 and CD8 T cell discrimination. Memory and naïve T cells were gated, followed by discrimination of $CXCR3^-$ and $CXCR3^+$ cells, and bonafide TSCM cells ($CD27^+CD95^+$) were gated from the $CXCR3^-$ fraction.

Table S1. nPOD donor and corresponding experiment information.

| case_id | RR_id | donor_type | experiment |
|---|---|---|---|
| 6374 | SAMN15879427 | No Diabetes | scRNAseq, CyTOF |
| 6131 | SAMN15879188 | No Diabetes | Fresh flow cytometry |
| 6174 | SAMN15879230 | No Diabetes | CyTOF, flow confirmation |
| 6178 | SAMN15879234 | No Diabetes | CyTOF |
| 6212 | SAMN15879268 | Type 1 Diabetes | CyTOF, flow confirmation |
| 6228 | SAMN15879284 | Type 1 Diabetes | CyTOF |
| 6238 | SAMN15879294 | No Diabetes | CyTOF |
| 6243 | SAMN15879299 | Type 1 Diabetes | CyTOF |
| 6247 | SAMN15879303 | Type 1 Diabetes | CyTOF |
| 6264 | SAMN15879318 | Type 1 Diabetes | CyTOF, flow confirmation |
| 6266 | SAMN15879320 | Type 1 Diabetes | CyTOF |
| 6282 | SAMN15879336 | No Diabetes | CyTOF |
| 6306 | SAMN15879360 | Type 1 Diabetes | CyTOF |
| 6317 | SAMN15879371 | No Diabetes | CyTOF |
| 6330 | SAMN15879384 | Type 1 Diabetes | Fresh flow cytometry |
| 6336 | SAMN15879390 | No Diabetes | CyTOF, flow confirmation |
| 6341 | SAMN15879395 | Type 1 Diabetes | CyTOF |
| 6371 | SAMN15879424 | Type 1 Diabetes | CyTOF, flow confirmation |
| 6375 | SAMN15879428 | No Diabetes | CyTOF |
| 6380 | SAMN15879433 | Type 1 Diabetes | Fresh flow cytometry |
| 6384 | SAMN15879437 | No Diabetes | Fresh flow cytometry |
| 6385 | SAMN15879438 | No Diabetes | Fresh flow cytometry |
| 6386 | SAMN15879439 | No Diabetes | CyTOF |
| 6387 | SAMN15879440 | No Diabetes | CyTOF, fresh flow cytometry, flow confirmation |
| 6389 | SAMN15879442 | No Diabetes | Fresh flow cytometry |
| 6391 | SAMN15879444 | No Diabetes | CyTOF |
| 6393 | SAMN15879446 | No Diabetes | Fresh flow cytometry |
| 6397 | SAMN15879450 | Autoantibody Positive | Fresh flow cytometry |
| 6400 | SAMN15879453 | Autoantibody Positive | Fresh flow cytometry |
| 6410 | SAMN15879463 | Other - No Diabetes | Fresh flow cytometry |
| 6412 | SAMN15879465 | No Diabetes | Fresh flow cytometry |
| 6413 | SAMN15879466 | No Diabetes | CyTOF,Fresh flow cytometry, flow confirmation |
| 6414 | SAMN15879467 | Type 1 Diabetes | scRNAseq, CyTOF, Fresh flow cytometry |

| | | | |
|---|---|---|---|
| 6415 | SAMN15879468 | No Diabetes | Fresh flow cytometry |
| 6417 | SAMN15879470 | No Diabetes | Fresh flow cytometry |
| 6418 | SAMN15879471 | Type 1 Diabetes | scRNAseq |
| 6420 | SAMN15879473 | No Diabetes | Fresh flow cytometry |
| 6422 | SAMN15879475 | Type 1 Diabetes | Fresh flow cytometry |
| 6424 | SAMN15879477 | Autoantibody Positive | Fresh flow cytometry |
| 6425 | SAMN15879478 | No Diabetes | Fresh flow cytometry |
| 6429 | SAMN15879482 | Autoantibody Positive | Fresh flow cytometry |
| 6432 | SAMN15879485 | Type 1 Diabetes | Fresh flow cytometry |
| 6440 | SAMN15879493 | No Diabetes | Fresh flow cytometry |
| 6457 | SAMN15879510 | Type 1 Diabetes | scRNAseq |
| 6461 | SAMN15879514 | No Diabetes | scRNAseq |
| 6462 | SAMN15879515 | No Diabetes | Flow confirmation |
| 6482 | SAMN15879535 | No Diabetes | scRNAseq |
| 6493 | SAMN15879546 | No Diabetes | scRNAseq |
| 6536 | SAMN18242780 | Type 1 Diabetes | scRNAseq |
| 6548 | SAMN25652259 | No Diabetes | scRNAseq |
| 6550 | SAMN25652261 | Type 1 Diabetes | scRNAseq |
| 6551 | SAMN25652262 | Type 1 Diabetes | scRNAseq |
| 6566 | SAMN33284286 | Type 1 Diabetes | scRNAseq |
| 6578 | SAMN33284295 | Type 1 Diabetes | scRNAseq, Flow confirmation |
| 6579 | SAMN33284296 | Type 1 Diabetes | scRNAseq |
| 6586 | SAMN38117306 | No Diabetes | scRNAseq |
| 6606 | SAMN40555593 | Type 1 Diabetes | Flow confirmation |
| 6610 | SAMN44486496 | Type 1 Diabetes | scRNAseq |
| 6614 | SAMN44486500 | No Diabetes | scRNAseq |

Table S2. Flow cytometry panel examining T cell phenotype in fresh tissue.

| Antigen | Fluorophore | Clone | Manufacturer |
|---|---|---|---|
| CCR7 | BV421 | G043H7 | Biolegend |
| CD4 | FITC | RPA-T4 | BioLegend |
| CD45RA | PerCP-Cy5.5 | HI100 | BioLegend |
| CXCR5 | PE | J252D4 | BioLegend |
| CXCR3 | PE-Cy7 | G025H7 | BioLegend |
| PD-1 | APC | eBioJ105 | eBioscience |
| CD3 | APC-H7 | SK7 | BD |
| Live/Dead | Live Dead Yellow | NA | Invitrogen |

Table S3. Flow cytometry panel assessing stem-like and exhausted T cell features in pLN.

| Marker | Fluorophore | Catalog | Vendor |
|---|---|---|---|
| Zombie Viability Die | Aqua | 423101 | Biolegend |
| CD45 | Spark blue 574 | 368558 | Biolegend |
| CD3 | Spark YG593 | 344868 | Biolegend |
| CD4 | Spark UV387 | 344686 | Biolegend |
| CD8 | BUV805 | 568334 | BD/Fisher |
| CD45RA | BUV563 | 612926 | BD/Fisher |
| CD45RO | R718 | 567536 | BD/Fisher |
| CD127 | BV711 | 351328 | Biolegend |
| PD1 | BV605 | 329924 | Biolegend |
| CXCR3 | BV421 | 353716 | Biolegend |
| CXCR5 | AF647 | 356906 | Biolegend |

| | | | |
|---|---|---|---|
| CD95 | RB780 | 569126 | BD/Fisher |
| CD27 | BUV496 | 751678 | BD/Fisher |
| CD57 | BV785 | 393330 | Biolegend |
| CD69 | RB705 | 570278 | BD/Fisher |
| CD28 | BUV395 | 742530 | BD/Fisher |
| CD5 | RY703 | 771019 | BD/Fisher |
| CD122 | BB515 | 566059 | BD |
| TCF1 | PE | 655207 | Biolegend |
| TOX | AF647 | 568356 | BD |

Table S4. Flow cytometry panel examining pLN $CD8^+$ T cell functionality.

| Marker | Fluorophore | Catalog | Vendor |
|---|---|---|---|
| Zombie Viability Die | Aqua | 423101 | Biolegend |
| CD3 | Spark YG593 | 344868 | Biolegend |
| CD4 | Spark UV387 | 344686 | Biolegend |
| CD8 | APC | 301014 | Biolegend |
| CD45RA | BUV563 | 612926 | BD/Fisher |
| CD45RO | R718 | 567536 | BD/Fisher |
| CXCR3 | BV421 | 353716 | Biolegend |
| CD95 | RB780 | 569126 | BD/Fisher |
| CD27 | BUV496 | 751678 | BD/Fisher |
| CD28 | BUV395 | 742530 | BD/Fisher |
| IFNG | PeCy7 | 502528 | Biolegend |
| TNFa | BV650 | 502938 | Biolegend |
| IL2 | BV750 | 500359 | Biolegend |

Table S5. Mass cytometry panel.

| Antigen | Isotope |
|---|---|
| CD57 | In113Di |
| CD19 | Nd142Di |
| CD4 | Nd143Di |
| CD8 | Nd144Di |
| CD16 | Nd145Di |
| IgD | Nd146Di |
| CD20 | Sm147Di |
| CD14 | Nd148Di |
| CD56 | Sm149Di |
| CD3 | Nd150Di |
| CD103 | Eu151Di |
| TCRgd | Sm152Di |
| CD45RA | Eu153Di |
| TIGIT | Sm154Di |
| CCR6 | Gd155Di |
| CXCR3 | Gd156Di |
| CD86 | Gd157Di |
| CD27 | Gd158Di |
| CD11c | Tb159Di |
| CCR7 | Gd160Di |
| CD69 | Dy162Di |
| CXCR5 | Dy164Di |
| CD127 | Ho165Di |
| CD33 | Er166Di |
| CD28 | Er167Di |
| CD24 | Er168Di |
| ICOS | Tm169Di |
| CD161 | Er170Di |
| CD226 | Yb171Di |
| CD38 | Yb172Di |
| CD123 | Yb173Di |
| PD1 | Yb174Di |
| HLADR | Lu175Di |
| CD25 | Yb176Di |
| CD11b | Bi209Di |

Table S6. TotalSeq antibodies and dextramer information

| Antibody/Dextramer | Item No | Dextramer Category |
|---|---|---|
| anti-human CD4 | 344651 | |
| anti-human CD8 | 344753 | |
| anti-human CD69 | 310951 | |
| anti-human CD27 | 302853 | |
| anti-human CD45RA | 304163 | |
| anti-human CD25 | 302649 | |
| anti-human CD127 (IL-7Rα) | 351356 | |
| HLA-A 0201 NLVPMVATV | | CMV |
| HLA-A 0201 VLEETSVML | | CMV |
| HLA-A 0201 ILGFVFTLTV | | MP |
| HLA-A 0201 KLGEFYNQMM | | Influenza |
| DRB1 0401 PKYVKQNTLKLAT | | Influenza |
| HLA-A 0101 VLFGLGFAI | | IGRP |
| HLA-A 0101 HLVEALYLV | | Insulin |
| HLA-A 0101 ALWGPDPAAA | | Insulin |
| HLA-A 0101 VMNILLQYVV | | GAD65 |
| HLA-A 0101 MVWESGCTV | | IA2 |
| DRB1*0301 IAFTSEHSHFSLK | | GAD65 |
| DRB1*0301 NFIRMVISNPAAT | | GAD65 |
| DRB1*0301 GAGSLQPLALEGSLQKRG | | Proinsulin |
| HLA-B*0801 AAKGRGAAL | | General negative control |
| HLA-A*0201 ALIAPVHAV | | |
| DRB1*0401 PVSKMRMATPLLMQA | | CLIP peptide |

Table S7. Phenocycler panel information

| | **Atto 550** | | | **Alexa Fluor 647** | | | **Alexa Fluor 750** | | |
|---|---|---|---|---|---|---|---|---|---|
| **Cycle Order** | **Target** | **Barcode** | **Exposure Time (ms)** | **Target** | **Barcode** | **Exposure Time (ms)** | **Target** | **Barcode** | **Exposure Time (ms)** |
| 1 | IAPP | BX032 | 150 | CD4 | BX003 | 150 | CD107a | BX006 | 150 |
| 2 | CD31 | BX001 | 125 | CD68 | BX015 | 100 | SMA | BX013 | 100 |
| 3 | CD44 | BX005 | 150 | CD99 | BX010 | 75 | Vimentin | BX022 | 100 |
| 4 | ECAD | BX014 | 150 | CD66 | BX016 | 150 | IDO1 | BX027 | 150 |
| 5 | INS | BX002 | 100 | CD11c | BX024 | 60 | Ker8-18 | BX107 | 150 |

| | | | | | | | | | |
|---|---|---|---|---|---|---|---|---|---|
| 6 | CD38 | BX089 | 150 | CD34 | BX025 | 150 | HLADR | BX033 | 150 |
| 7 | iNOS | BX023 | 100 | B3Tub | BX017 | 150 | M2Gal3 | BX035 | 80 |
| 8 | CD8 | BX026 | 150 | FOXP3 | BX031 | 150 | PCNA | BX036 | 150 |
| 9 | CD57 | BX028 | 80 | GranB | BX041 | 150 | Ki67 | BX047 | 150 |
| 10 | HLAA | BX029 | 100 | Col_IV | BX042 | 150 | CD20 | BX064 | 150 |
| 11 | VISTA | BX040 | 150 | PD-1 | BX046 | 150 | PanCK | BX066 | 75 |
| 12 | LAG3 | BX055 | 125 | TCF-1 | BX061 | 150 | STT | BX037 | 150 |
| 13 | TOX | BX060 | 150 | ICOS | BX065 | 150 | Caveolin | BX086 | 150 |
| 14 | CD163 | BX069 | 125 | PD-L1 | BX067 | 150 | EpCAM | BX091 | 150 |
| 15 | CD79a | BX090 | 125 | CD3e | BX080 | 150 | Ker5 | BX101 | 125 |
| 16 | GCG | BX020 | 75 | Bcl-2 | BX085 | 125 | BetaActin | BX117 | 100 |
| 17 | MPO | BX098 | 125 | CD56 | BX088 | 150 | - | | - |
| 18 | CD39 | BX099 | 150 | Iba1 | BX021 | 100 | - | | - |
| 19 | SOX2 | BX102 | 150 | CD209 | BX329 | 150 | - | | - |
| 20 | PDPN | BX121 | 150 | CD11b | BX524 | 80 | - | | - |
| 21 | CD206 | BX361 | 150 | TP63 | BX093 | 100 | - | | - |
| 22 | - | - | - | - | - | - | - | | - |
| 23 | - | - | - | - | - | - | - | | - |
| 24 | - | - | - | - | - | - | - | | - |

Table S8. Demographic information for mass cytometry (CyTOF) cohort.

| Clinical Status | n | % Male | Age (yr) | % AAB+ | Disease Duration (yr) | % Insulitis + | % High Risk HLA |
|---|---|---|---|---|---|---|---|
| T1D | 10 | 80 | 19.5(18) | 90 | 5(23) | 90 | 100 |
| ND | 12 | 66.7 | 14.65(18.6) | 0 | NA | 0 | 66.7 |

Age and disease duration represented as median(range)

*High Risk HLA was defined by possession of risk associated alleles:
DRB1*0401,0402,0403,0404;DRB1*0301

Table S9. Demographic information for flow cytometry validation cohort.

| Clinical Status | n | % Male | Age (yr) | # AAB | Disease Duration (yr) | % Insulitis + | % High Risk HLA |
|---|---|---|---|---|---|---|---|
| T1D | 5 | 60 | 19.2±5.0 | 20% negative, 20% single, 20% double, 40% triple | 10.5±9.8 | 80 | 100 |
| AAB+ | 4 | 75 | 21.5±3.1 | 50% single, 50% double | NA | 25 | 75 |
| ND | 14 | 64 | 17.1±7.3 | | NA | 0 | 60 |

*High Risk HLA was defined by possession of risk associated alleles:
DRB1*0401,0402,0403,0404,0407;DRB1*0301

Table S10. Demographic information for scRNAseq cohort.

| Clinical Status | n | % Male | Age (yr) | % AAB+ | Disease Duration (yr) | % Insulitis+ | % High Risk HLA |
|---|---|---|---|---|---|---|---|
| T1D | 9 | 55 | 17.0±6.5 | 100 | 1.80±3.4 | 100 | 100 |
| ND | 7 | 71.4 | 17.1±3.1 | 0 | NA | 0 | 80 |

*High Risk HLA was defined by possession of risk associated alleles:
DRB1*0401,0402,0403,0404;DRB1*0301

Table S11. Differentially expressed genes between T1D (positive fold change) and ND (negative fold change) in T cell clusters (see supplemental values).
